# BAF complexes cooperate with lineage-restricted transcription factors to establish cell-state-specific regulatory programs

**DOI:** 10.64898/2026.11.08.764927

**Authors:** Jiayou He, Carla Ríos Arceo, Nadia Anderson, Willem Teunissen, Irene Paassen, Siem Jongsma, Elzo de Wit, Marcel Kool, Ning Qing Liu, Jarno Drost

## Abstract

BRG1/BRM-associated factor (BAF) complexes are essential regulators of chromatin organization and cell identity. Genes encoding BAF subunits are among the most frequently mutated in human cancer. Yet, how disruption of a single subunit gives rise to distinct tumor states remains poorly understood. Atypical teratoid rhabdoid tumors (ATRTs) provide a compelling model, as they are uniformly driven by bi-allelic loss of the BAF subunit *SMARCB1*, yet segregate into distinct molecular subtypes. Using integrated multi-omics profiling of patient-derived ATRT organoids and primary tumor tissues, we show that transcriptional and chromatin accessibility changes upon *SMARCB1* reconstitution are directed by subtype-and lineage-specific transcription factor (TF) repertoires. Genome-wide binding and transcriptomic analyses reveal that *SMARCB1* reconstitution enables the lineage-specific TF RFX4 to engage regulatory elements and activate neuronal differentiation programs, partially reprogramming ATRT-MYC toward an ATRT-SHH-like identity. Together, these findings demonstrate that *SMARCB1* loss disrupts TF-dependent chromatin priming, revealing that the lineage-specific regulatory output of BAF complexes is constrained by the underlying TF repertoire.

## Main

Approximately 20-25% of human cancers harbor alterations in genes encoding subunits of the BRG1/BRM-associated factor (BAF) complex^1,2^. BAF complexes are ATP-dependent chromatin remodelers that regulate gene expression by repositioning nucleosomes, thereby modulating chromatin accessibility^3^. They assemble into three distinct variants namely canonical BAF (cBAF), polybromo-associated BAF (pBAF), and non-canonical BAF (ncBAF), each defined by a unique subunit composition that confers specialized regulatory functions and genomic targeting^4–6^. Importantly, the composition of BAF complexes varies across different cell types and developmental stages, enabling them to regulate lineage-specific transcriptional programs^7–9^. Disruption of these context-dependent epigenetic mechanisms is increasingly recognized as a key driver of tumorigenesis^10^. However, how mutations in BAF complex components alter lineage identity and promote malignant transformation remains poorly understood.

A major challenge in studying the oncogenic consequences of BAF complex mutations is the genetic complexity of most adult malignancies, which are generally characterized by highly complex mutational landscapes. This complexity makes it difficult to disentangle the specific contribution of BAF complex dysfunction from the effects of cooperating genetic alterations. In contrast, pediatric cancers typically harbor fewer driver mutations and several aggressive pediatric tumor types are even driven by inactivation of a single BAF complex component^11,12^. These cancers therefore serve as tractable model systems for studying the role of BAF-mediated chromatin dysregulation in tumorigenesis. Malignant rhabdoid tumors (MRTs) represent a cancer type driven by mutation of a single BAF complex component. These highly aggressive pediatric malignancies are characterized by bi-allelic inactivation of the BAF subunit *SMARCB1* (also known as INI1/SNF5/BAF47) in approximately 95% of cases, whereas the remaining tumors harbor loss of *SMARCA4*^13–16^. MRTs primarily affect infants and can arise both intracranially as atypical teratoid rhabdoid tumors (ATRTs) and at extracranial sites^17,18^. Despite their remarkably simple genetic landscape, MRTs exhibit substantial phenotypic and molecular heterogeneity. In ATRTs, integrated transcriptomic and epigenomic analyses have identified three major molecular subtypes, sonic hedgehog (ATRT-SHH), MYC (ATRT-MYC), and tyrosinase (ATRT-TYR), each characterized by distinct gene regulatory programs and clinical behaviors^15,19^. The striking molecular diversity of ATRTs despite a common initiating genetic lesion raises the question of how loss of a single chromatin-remodeling complex component drives distinct lineage-specific oncogenic programs. Accumulating evidence suggests that ATRTs originate from developmental neuronal cells as a consequence of impaired differentiation and lineage specification during early development^20,21^. Previous studies by us and others have demonstrated that the tumorigenic consequences of *SMARCB1* loss depend on a specific developmental context and require additional, most likely non-genetic (e.g., epigenetic) events^22,23^. Consistent with this concept, reconstitution of *SMARCB1* was shown to reconfigure enhancer-promoter interactions, including MRT-specific looping of distal enhancers to the *MYC* promoter, highlighting the profound influence of BAF complex integrity on gene regulation^24–26^. Despite these advances, how *SMARCB1* loss disrupts lineage-specific regulatory networks to drive tumorigenesis remains underexplored.

Here, we use patient-derived ATRT organoids and primary tumor tissues to investigate how loss of the BAF complex subunit SMARCB1 rewires lineage-specific regulatory networks and locks cells into an undifferentiated cellular state. Our study shows that the developmental consequences of *SMARCB1* loss are dictated by the underlying transcription factor (TF) repertoire, providing a mechanistic framework for how disruption of a single BAF complex subunit gives rise to distinct tumor identities.

## Results

### *SMARCB1* reconstitution in ATRT organoids induces a growth arrest

To examine the functional consequences of restoring cBAF complex function in ATRT^27^, we reconstituted *SMARCB1* expression in multiple independent patient-derived ATRT organoids of different subtypes (n=3 ATRT-SHH; n=4 ATRT-MYC). As such, we lentivirally transduced ATRT organoids with either a luciferase gene (Control) or the wild-type *SMARCB1* gene (SMARCB1+) (Fig. 1a). *SMARCB1* reconstitution was confirmed by immunohistochemistry and western blot analysis (Fig. 1b,d, Extended Data Fig. 1a,b). Across all the models, irrespective of molecular subtype, *SMARCB1* reconstitution led to a robust proliferation arrest (Fig. 1e). In contrast to extracranial MRTs (ecMRTs)^22^, no notable morphological changes were observed upon *SMARCB1* re-expression in the majority of ATRT organoid cultures (Fig. 1c and Extended Data Fig. 1c). One notable exception was AT-MYC23, which underwent a morphological transition from a compact, grape-like spheroid architecture to a structure bearing neuron-like protrusions upon *SMARCB1* re-expression (Extended Data Fig. 1c). Together, these findings demonstrate that *SMARCB1* expression can be efficiently reconstituted in ATRT organoids, resulting in a proliferation arrest across molecularly distinct ATRT subtypes.

**Figure 1:**
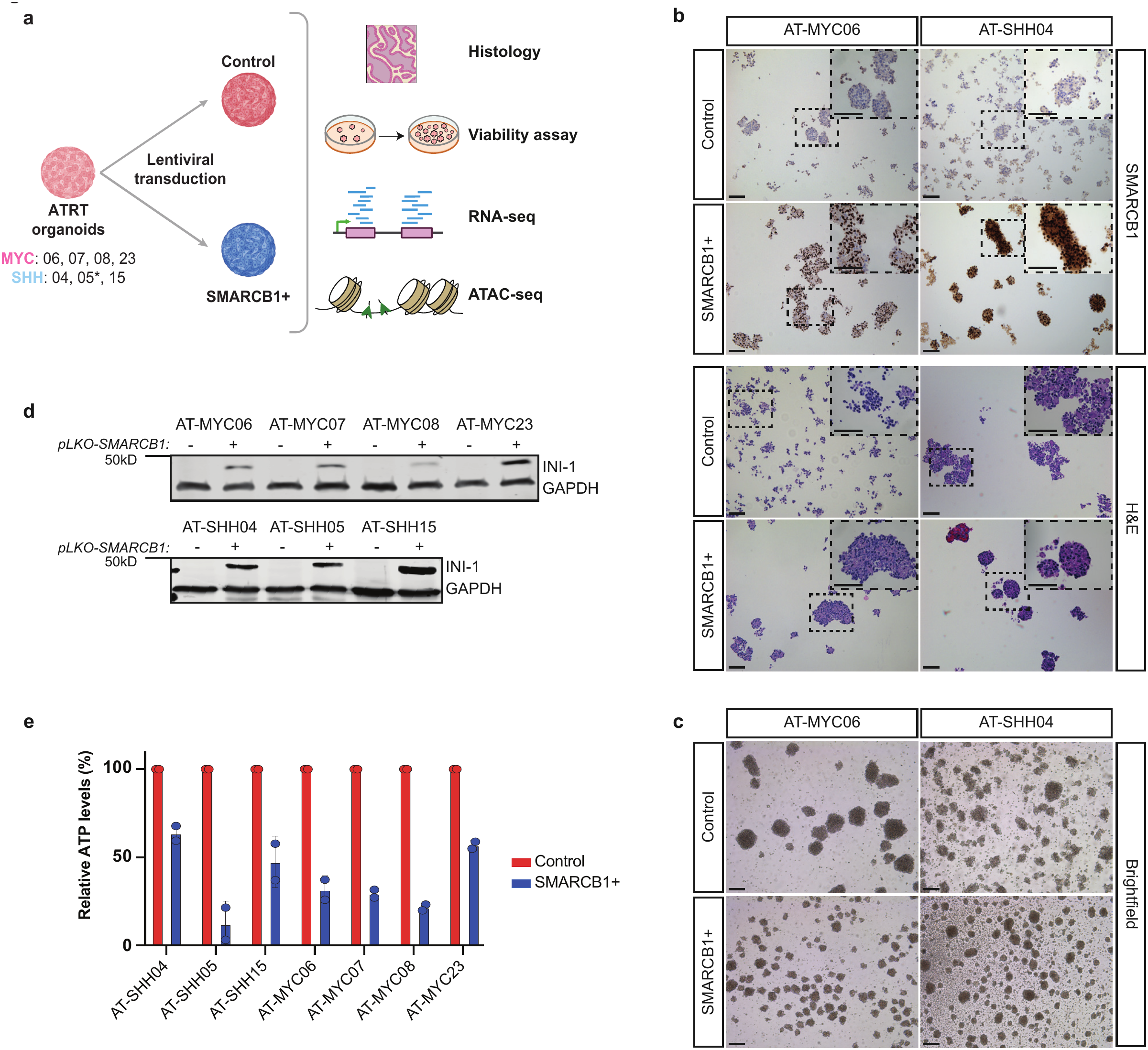
*SMARCB1* reconstitution in ATRT organoids induces a growth arrest. **a,** Schematic representation of the experimental design. * AT-SHH05 was excluded from ATAC-seq analyses due to technical limitations associated with reduced viability of this model following *SMARCB1* reconstitution. **b,** Representative images of SMARCB1 (INI-1) (top) and hematoxylin and eosin (H&E) (bottom) stainings on the indicated ATRT organoid models transduced with either a control (Control) or *SMARCB1* (SMARCB1+) expression plasmid. Scale bars = 100 μm. **c,** Brightfield images of ATRT organoid models with (SMARCB1+) and without (Control) *SMARCB1* reconstitution. Scale bars = 100 μm. **d,** Western blot of SMARCB1 expression in the indicated ATRT organoid models without (-) and with (+) *SMARCB1* reconstitution. GAPDH was used as loading control. **e,** Bar plot with relative ATP levels (%), measured by CellTiter-Glo (CTG), as a proxy for organoid proliferation five days after replating. Data are presented relative to the corresponding Control condition. Dots represent n=2 biological replicates.

### *SMARCB1* reconstitution in ATRT organoids induces subtype-specific gene expression changes without affecting subtype identity

To define SMARCB1-dependent, subtype-specific transcriptional programs in ATRT, we performed bulk RNA-sequencing (RNA-seq) on Control and SMARCB1+ ATRT-SHH (n=3) and ATRT-MYC (n=4) organoid models. As expected, *SMARCB1* transcript levels were significantly increased across all isogenic organoid pairs following *SMARCB1* reconstitution (Extended Data Fig. 2a). Unsupervised clustering revealed that samples segregated primarily according to molecular subtype (ATRT-MYC versus ATRT-SHH), with SMARCB1 status (Control versus SMARCB1+) representing the second major source of variation (Fig. 2a). Similarly, principal component analysis demonstrated a consistent global transcriptomic response to *SMARCB1* reconstitution across all organoid models (Extended Data Fig. 2b). Differential gene expression analysis of each isogenic organoid pair showed that *SMARCB1* reconstitution preferentially induced gene activation rather than repression (Extended Data Fig. 2c), consistent with SMARCB1 functioning primarily as a transcriptional activator^28^. Comparison of Control ATRT-SHH and ATRT-MYC organoids identified robust subtype-specific gene expression signatures. In particular, ATRT-MYC exhibited higher expression of multiple transcription factor (TF) families, including HOXA/B/C/D and FOX, whereas ATRT-SHH preferentially expressed TFs such as MYCL/N, OLIG1/2, RFX4, and SOX (Fig. 2b, left), consistent with earlier reports using ATRT patient samples^15,19,29^. Notably, more than 50% of these subtype-specific genes, including most TFs, retained their subtype preference following *SMARCB1* reconstitution (Fig. 2b, right), indicating that reconstitution of *SMARCB1* alone is insufficient to erase the core transcriptomic identity of each subtype.

**Figure 2:**
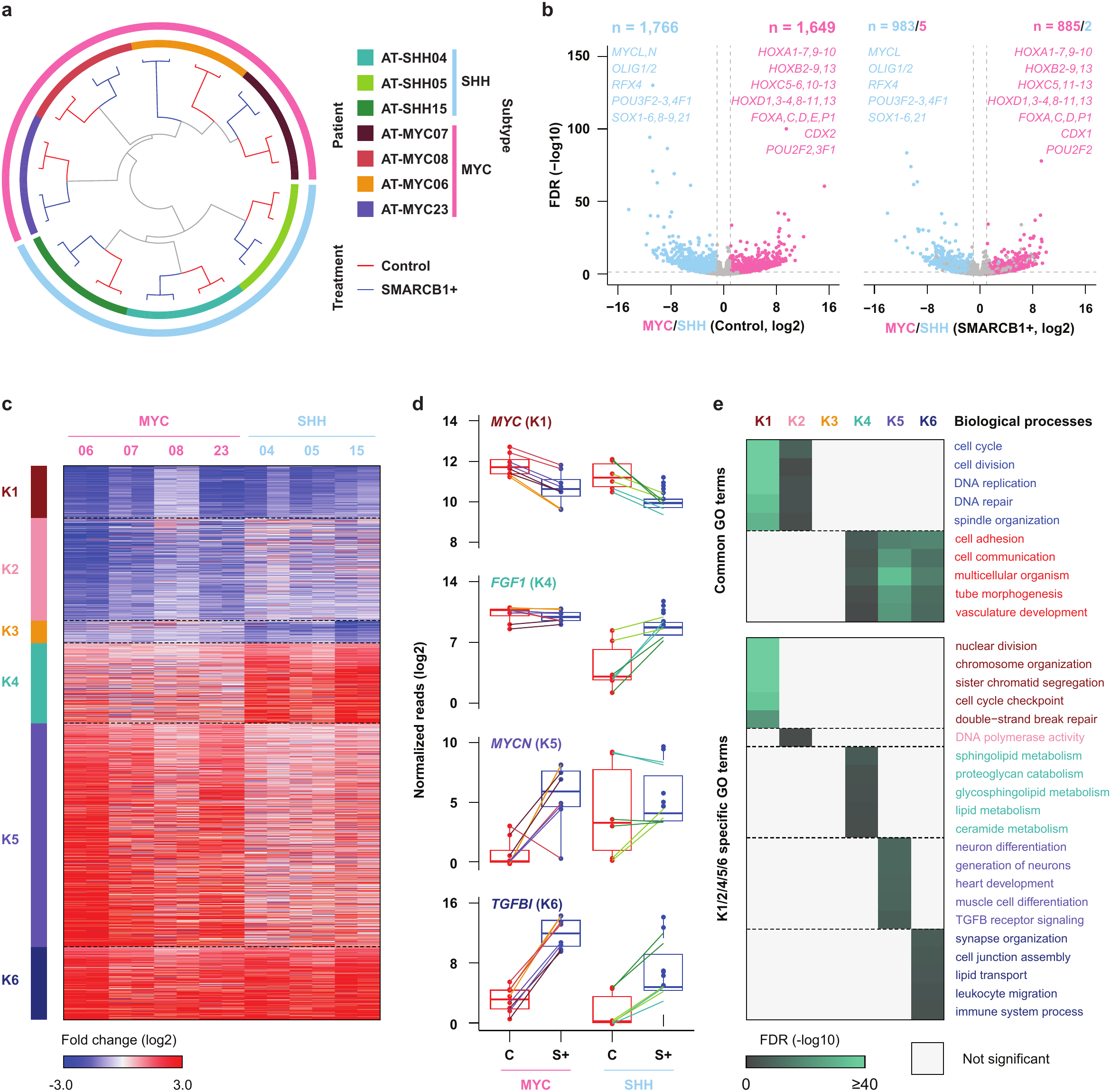
*SMARCB1* reconstitution in ATRT activates subtype-specific transcriptomic programs in ATRT. **a,** Circos plot showing clustering of seven ATRT organoid pairs based on transcriptomic profiles in the absence (Control) and presence (SMARCB1+) of *SMARCB1*. **b,** Differentially expressed genes between Control ATRT-SHH and Control ATRT-MYC organoids were identified and visualized as volcano plots using RNA-seq data from Control (left panel) and SMARCB1+ (right panel) conditions. Representative subtype-defining transcription factors are indicated and highlighted in pale violet red (MYC-specific) or sky blue (SHH-specific). **c,** Heatmap showing common (K1 and K6) and subtype-specific (K2-5) transcriptomic changes following *SMARCB1* reconstitution across ATRT organoid subtypes. **d,** Box plots showing representative common and subtype-specific genes affected by *SMARCB1* reconstitution in MYC and SHH ATRT organoids. Genes correspond to K-means clusters: *MYC* (K1), *FGF1* (K4), *MYCN* (K5), and *TGFBI* (K6). (C) Control; S+, SMARCB1+. **e,** Gene Ontology analysis of the six gene clusters highlighting enriched biological processes. Common enrichments are shared across multiple clusters (at least two), whereas subtype-specific enrichments are indicated for K1, K2, K4, K5, and K6. The scale bar represents −log10 false discovery rate (FDR).

To distinguish shared from subtype-specific transcriptomic responses to *SMARCB1* reconstitution, we next compared Control and SMARCB1+ organoids across both subtypes (Fig. 2c). Nearly all differentially expressed genes (2,595 out of 2,598) could be assigned to one of six expression clusters. Clusters K1 and K6 contained genes consistently downregulated or upregulated, respectively, in both ATRT subtypes following *SMARCB1* reconstitution, including *MYC* (K1) and *TGFBI* (K6) (Fig. 2d). Gene Ontology (GO) analysis revealed that K1 was highly enriched for cell cycle, DNA replication, and chromosome segregation genes (Fig. 2e), consistent with the proliferation arrest induced by *SMARCB1* reconstitution (Fig. 1e and Extended Data Fig. 1d). In contrast, K6 was enriched for genes involved in cell communication, tube morphogenesis, and synapse organization (Fig. 2e).

In addition to these shared responses, we identified subtype-specific transcriptomic changes. Clusters K2 and K3 comprised genes selectively downregulated following *SMARCB1* reconstitution in ATRT-MYC and ATRT-SHH, respectively (Fig. 2c). Although no significant GO terms were identified for K3 because of its limited size, K2 was enriched for cell cycle-and DNA replication-associated genes (Fig. 2e). Conversely, clusters K4 and K5 contained genes preferentially induced in ATRT-SHH and ATRT-MYC, respectively, including *FGF1* (K4) and *MYCN* (K5) (Fig. 2c,d). GO analysis showed that K4 was enriched for lipid metabolic processes, whereas K5 was associated with neural differentiation, muscle differentiation, and heart development (Fig. 2e).

Interestingly, genes in K4 and K5 reached comparable expression levels in both subtypes following *SMARCB1* reconstitution but were selectively silenced in Control ATRT-SHH and ATRT-MYC organoids, respectively (Fig. 2d and Extended Data Fig. 2d). Thus, in the *SMARCB1*-deficient state, K4 genes are specifically suppressed in ATRT-SHH, whereas K5 genes are selectively suppressed in ATRT-MYC (Extended Data Fig. 2e). In contrast, genes in the commonly affected clusters (K1 and K6) displayed similar expression level between the subtypes in the *SMARCB1*-null condition and had similar degree of response to *SMARCB1* reconstitution.

Together, these findings indicate that *SMARCB1* reconstitution elicits a conserved transcriptional program associated with growth arrest and differentiation while preserving the core subtype-specific transcriptomic identities of ATRT-SHH and ATRT-MYC.

### SMARCB1-dependent chromatin remodeling is marked by subtype-specific transcription factors

SMARCB1 is an essential subunit of the cBAF and pBAF complexes that regulate gene expression by repositioning nucleosomes to establish accessible chromatin^30^. To define how *SMARCB1* loss alters the chromatin landscape, we profiled chromatin accessibility using assay for transposase-accessible chromatin using sequencing (ATAC-seq) in six ATRT organoid models (n=2 ATRT-SHH; n=4 ATRT-MYC) in the absence and presence of SMARCB1. In contrast to the transcriptomic analyses (Fig. 2a), unsupervised clustering and principal component analyses showed that SMARCB1 status, rather than molecular subtype, now was the primary determinant of chromatin accessibility (Fig. 3a and Extended Data Fig. 3a). These findings suggest that *SMARCB1* reconstitution re-establishes a core BAF-dependent chromatin accessibility landscape shared across ATRT subtypes.

**Figure 3:**
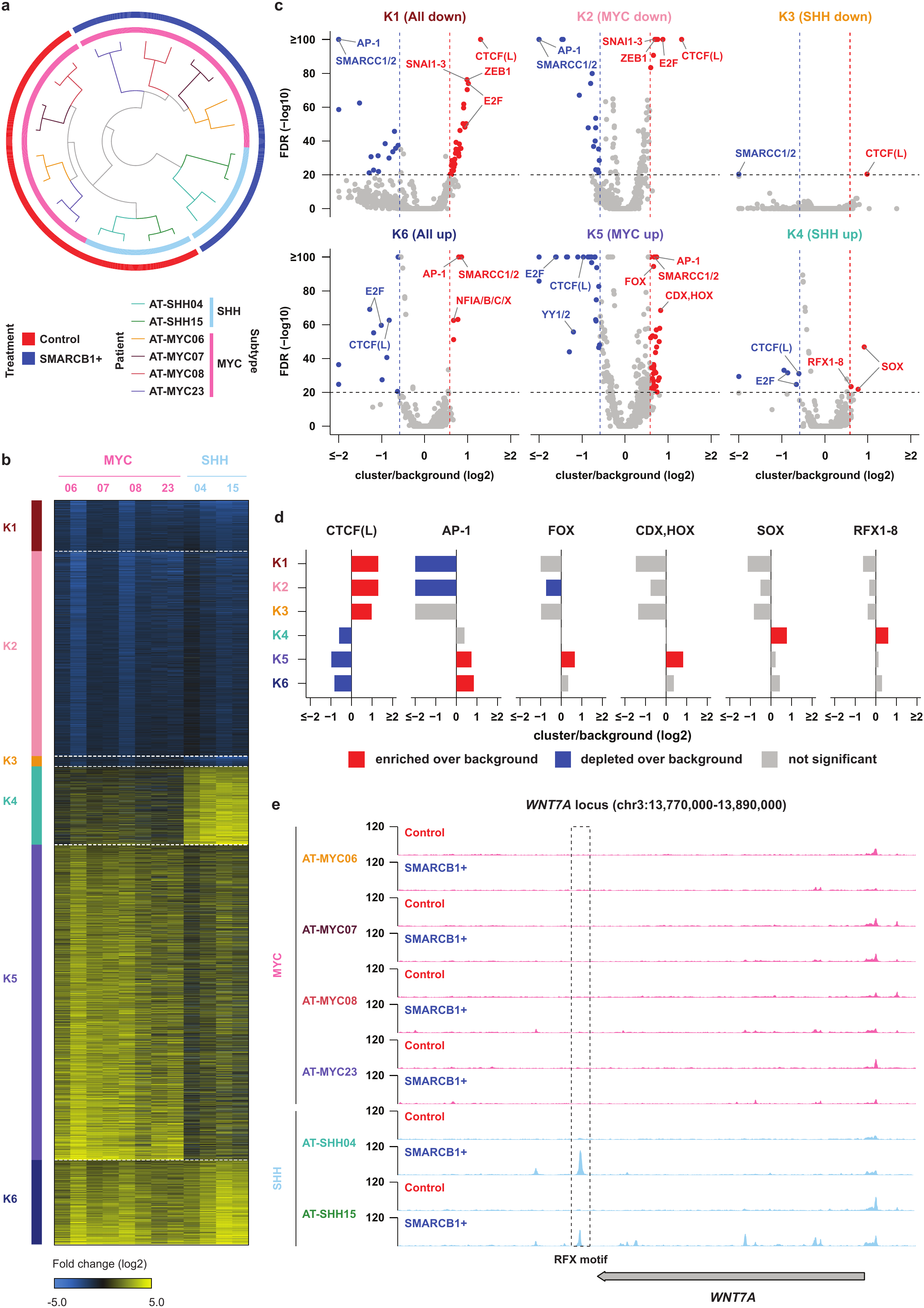
SMARCB1-dependent chromatin remodeling is marked by subtype-specific transcription factors. a, Circos plot showing clustering of six ATRT organoid models based on genome-wide chromatin accessibility in the absence (Control) and presence (SMARCB1+) of *SMARCB1*. **b,** Heatmap depicting six classes of differentially accessible chromatin regions (K1–K6), comprising common (K1, K6) and subtype-specific (K2–K5) responses to *SMARCB1* reconstitution across ATRT organoid models. **c,** Volcano plots showing TF binding motif enrichment across the six chromatin accessibility clusters. Each dot represents a TF motif. Positive log2 fold enrichment (x-axis) indicates overrepresentation in the indicated cluster relative to the background (all accessible regions), whereas negative values indicate depletion. **d,** Bar plots showing enrichment and depletion of representative TF motifs across the six chromatin clusters (K1–K6). Bars indicate log2 fold change relative to background. Red indicates significant enrichment, blue significant depletion, and grey non-significant changes. **e,** Representative genomic locus of K4 chromatin regions near the *WNT7A* gene. The dashed box highlights a putative regulatory element containing an RFX motif that gains accessibility upon *SMARCB1* reconstitution in ATRT-SHH organoids.

Analysis of differential accessible regions (DARs) within each isogenic organoid pair (Control versus SMARCB1+) revealed substantial variability in the number of DARs across models (Extended Data Fig. 3b). Nevertheless, most of the models (four out of six) had considerably more regions gaining than losing accessibility following *SMARCB1* reconstitution (Extended Data Fig. 3b). Together with the RNA-seq analyses (Extended Data Fig. 2b,c), these data indicate that, at least in ATRTs, SMARCB1 primarily functions to promote chromatin accessibility and subsequently activate transcription.

To distinguish shared and subtype-specific chromatin responses, we compared chromatin accessibility between Control and SMARCB1+ organoids within each of the two molecular subtypes. Nearly all DARs (27,839 out of 27,840) could be assigned to one of six clusters: regions commonly losing (K1) or gaining (K6) accessibility after SMARCB1 restoration in both subtypes, regions selectively losing accessibility in ATRT-MYC (K2) or ATRT-SHH (K3), and regions selectively gaining accessibility in ATRT-SHH (K4) or ATRT-MYC (K5) (Fig. 3b). Genomic annotation showed that K1-K3 DARs were preferentially located at promoter-proximal regions, whereas K4-K6 DARs were predominantly associated with distal regulatory elements including many enhancers^31,32^ (Extended Data Fig. 3c). These findings are consistent with previous studies demonstrating that SMARCB1 primarily regulates chromatin accessibility at enhancer elements^24,33–35^.

Given that ATRT subtypes are thought to arise from distinct developmental lineages^20,36,37^, we reasoned that subtype-specific DARs might reflect lineage-restricted BAF complex activity. To identify TFs associated with these chromatin states, we performed motif enrichment analysis across the six DAR clusters. Consistent with previous findings in ecMRTs^22^, regions losing accessibility following *SMARCB1* reconstitution (K1) were enriched for CTCF motifs and depleted of SMARCC1/2 and AP-1 motifs (Fig. 3c,d), suggesting that SMARCB1 selectively reduces accessibility at CTCF-bound sites while restoring accessibility at enhancer-associated regions^24,38,39^. In addition, K1-K3 regions were enriched for E2F motifs (Fig. 3c), consistent with the proliferative program that is suppressed upon *SMARCB1* reconstitution. Genome-wide TF footprinting across individual organoid pairs confirmed these motif enrichment patterns (Extended Data Fig. 3d).

Analysis of the subtype-specific accessibility clusters identified distinct TF repertoires associated with each ATRT subtype. K5 DARs (MYC up) were enriched for consensus motifs of the FOX, CDX and HOX TF families (Fig. 3c,d), in agreement with previous studies^15,19,39^. The TFs in these three families are the key regulators of neurodevelopment in brain and spinal cord^40–42^ via their regulatory functions at lineage-specific enhancers^43–46^. In contrast, K4 DARs (SHH up) were enriched for motifs recognized by SOX and RFX family TFs (Fig. 3c,d). SOX factors have previously been implicated in neural lineage specification and ATRT-SHH biology^15,47,48^. The RFX family has been linked primarily to neural development and ciliogenesis and while transcriptional signals of this TF family have been previously described in ATRT^49–51^, it has not been functionally investigated.

To determine whether RFX factors are actively engaged at these regions, we examined Tn5 insertion patterns surrounding RFX motifs within K4 DARs. Pronounced RFX footprints were detected exclusively in ATRT-SHH organoids (AT-SHH04 and AT-SHH15) following *SMARCB1* reconstitution (Extended Data Fig. 3e), indicating SMARCB1-dependent occupancy of RFX-binding sites specifically in this subtype. These observations suggest that one or more RFX proteins interact with SMARCB1-containing BAF complexes to establish or maintain accessibility at SHH-specific regulatory elements (Fig. 3b,e). Notably, among all RFX family members, only *RFX4* was selectively expressed in ATRT-SHH organoids (Extended Data Fig. 3f), identifying RFX4 as the most likely mediator of the SHH-specific chromatin accessibility program induced upon *SMARCB1* reconstitution.

Together, these findings suggest that SMARCB1 acts as a critical coregulator of lineage-specific TFs to sustain an active state of subtype-specific regulatory elements. By restoring chromatin accessibility at lineage-specific binding sites, SMARCB1 may facilitate pre-existing TFs to engage their target loci, suggesting that BAF complex activity is constrained by the underlying TF repertoire.

### RFX4 is selectively active in ATRT-SHH and suppresses ATRT growth regardless of subtype

To extend our findings to primary tumors, we reanalyzed previously published paired single-nucleus RNA-and ATAC-seq datasets from MRTs^24,52^. In total, 28,798 nuclei from nine ecMRTs and eight ATRTs, including ATRT-TYR (n = 2), ATRT-SHH (n = 5), and ATRT-MYC (n = 1), were analyzed (Fig. 4a). We applied SCENIC+ to infer extended regulons (eRegulons), integrating TF expression, predicted target genes, and associated cis-regulatory elements to identify subtype-specific regulatory networks. Across all tumors, 97 active eRegulons were identified. Ranking eRegulons by activity revealed distinct subtype-specific regulatory programs (Fig. 4b). Consistent with previous studies, ATRT-TYR was enriched for the *OTX2* eRegulon, whereas ATRT-SHH exhibited strong *SOX2* eRegulon activity, reflecting activation of neural stem and progenitor-like transcriptional programs^19,53^. No ATRT-MYC-specific eRegulon was identified, likely owing to the inclusion of only a single ATRT-MYC sample.

**Figure 4:**
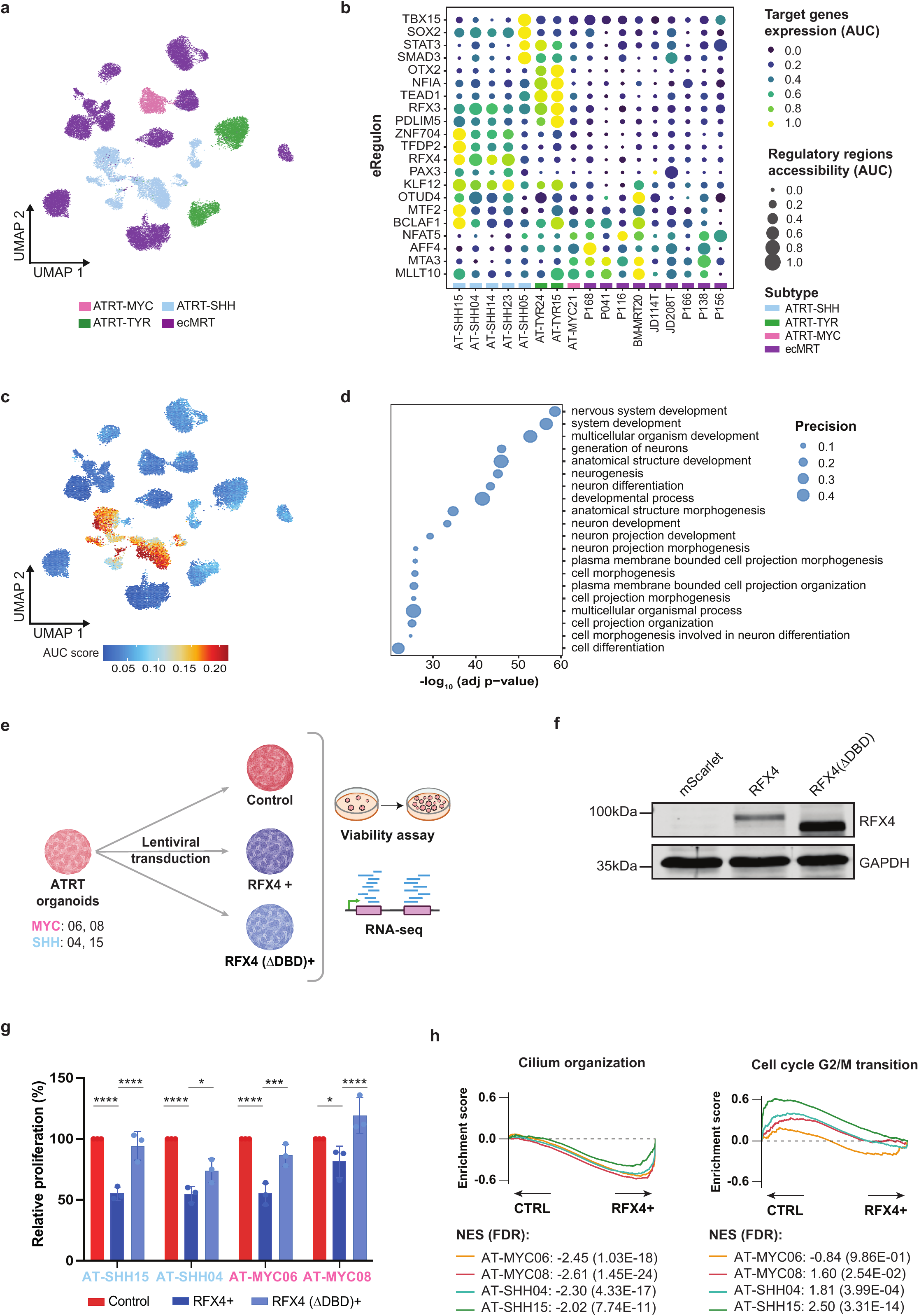
RFX4 is selectively active in ATRT-SHH and suppresses ATRT growth regardless of subtype. a, UMAP projection of snMultiome (RNA-seq + ATAC-seq) data from 17 primary MRT tissues, colored by molecular subtype. **b,** Dot plot showing genes comprising the top five “+/+” eRegulons identified for each ATRT subtype. eRegulon activity (gene AUC score) is displayed across all ATRT and ecMRT samples. “+/+” denotes a positive correlation between TF expression, accessibility of its associated regulatory regions, and expression of predicted target genes. **c,** UMAP projection of snMultiome data colored by RFX4 eRegulon activity (gene AUC score). **d,** Gene Ontology enrichment analysis showing the most significantly enriched biological processes in cells with high RFX4 eRegulon activity (“RFX4-high”). **e,** Schematic overview of the experimental design for ectopic expression of *RFX4* and the DNA-binding-deficient mutant *RFX4*(ΔDBD). ΔDBD, DNA-binding domain deletion. **f,** Western blot showing RFX4 protein expression in ATRT organoids in the indicated conditions. GAPDH was used as loading control. **g,** Bar plot with relative ATP levels (%), measured by CellTiter-Glo (CTG), as a proxy for organoid proliferation seven days after replating. Data are presented relative to the corresponding Control condition. Statistical significance was determined by two-way ANOVA followed by Tukey’s multiple-comparisons test. ****adjusted P < 0.0001; ***adjusted P < 0.001; **adjusted P < 0.01; *adjusted P < 0.05; ns, not significant. Dots represent n=3 biological replicates. **h,** Gene Set Enrichment Analysis (GSEA) of bulk RNA-seq data following ectopic *RFX4* expression (RFX4+ versus Control) showing significant enrichment of the Gene Ontology gene sets “Cilium organization” (left) and “Cell cycle G2/M transition” (right) across all four ATRT organoid models. Enrichment curves are shown for each organoid model individually. Normalized enrichment scores (NES) and false discovery rates (FDR) are indicated. All gene sets were significantly enriched (FDR < 10^-10^).

Importantly, *RFX4* emerged as one of the highest-ranked eRegulons specifically in ATRT-SHH tumors (Fig. 4b,c), whereas RFX3 activity was preferentially detected in ATRT-TYR (Extended Data Fig. 4b). Consistent with our organoid data, *RFX4* expression was restricted to ATRT-SHH cells (Extended Data Fig. 4a). Moreover, both *RFX4* expression and eRegulon activity exhibited a gradient across ATRT-SHH cells (Fig. 4c and Extended Data Fig. 4a), and cells with high *RFX4* activity were enriched for neuronal differentiation-associated gene programs (Fig. 4d and Extended Data Fig. 4c), consistent with a more differentiated cellular state. Together, these findings identify RFX4 as a candidate lineage-defining TF underlying the ATRT-SHH regulatory program in the absence of SMARCB1.

Given the prominent *RFX4* activity observed in SMARCB1-deficient ATRT-SHH tumors, we next asked whether RFX4 is sufficient to drive the SHH-specific transcriptional program. To this end, we ectopically expressed *RFX4* in two organoid models of each ATRT subtype (Fig. 4e,f). *RFX4* expression significantly reduced organoid growth irrespective of subtype (Fig. 4g), and this effect was abolished by deletion of the DNA-binding domain (RFX4ΔDBD), demonstrating that growth suppression depends on sequence-specific DNA binding. RNA-seq further showed that RFX4 elicited highly similar transcriptomic responses in both ATRT subtypes, characterized primarily by repression of cell cycle genes and induction of ciliogenesis associated gene signature (Fig. 4h and Extended Data Fig. 4d).

Together, these findings show that expression of *RFX4* induces a cell cycle arrest regardless of ATRT subtype and suggests that, in the absence of SMARCB1 and despite its activity in ATRT-SHH tumors, RFX4 alone is insufficient to establish an SHH-specific transcriptional program.

### SMARCB1 facilitates RFX4-dependent activation of the ATRT-SHH regulatory program

To determine whether SMARCB1 enables the lineage-specific functions of RFX4, we generated dual-transduced ATRT-MYC and ATRT-SHH organoids expressing SMARCB1 and/or RFX4 (Fig. 5a). Bulk RNA-seq revealed that molecular subtype remained the primary determinant of global transcriptional identity (Fig. 5b and Extended Data Fig. 5a). However, whereas ectopic *RFX4* expression alone induced only modest transcriptomic changes, co-expression of *SMARCB1* and *RFX4* elicited a substantially stronger transcriptional response in ATRT-MYC than *SMARCB1* reconstitution alone (Extended Data Fig. 5a,b). In contrast, *RFX4* had little additional effect in *SMARCB1*-reconstituted ATRT-SHH organoids, consistent with the higher endogenous expression of *RFX4* in this subtype (Extended Data Fig. 3f). Next, we specifically identified the differentially expressed genes (n=1,328) from the SMARCB1+/RFX4+ condition of AT-MYC08, because these gene expression changes of ATRT-MYC caused by the dual transduction may implicate transcriptomic adaption of this subtype toward ATRT-SHH. Notably, PCA using these selected genes showed that double-transduced ATRT-MYC organoids indeed shifted toward a more ATRT-SHH-like transcriptomic state (Fig. 5b). Consistent with this observation, combined *SMARCB1*/*RFX4* expression selectively induced the ATRT-SHH-associated K4 gene signature while further repressing the common proliferative K1 program in ATRT-MYC (Extended Data Fig. 5c). Furthermore, classification of the genes selected from AT-MYC08 dual transduction revealed coordinated repression of cell cycle and DNA replication genes (DT-K1 and DT-K2) together with activation of neuronal differentiation, cell adhesion, and synaptic signaling programs (DT-K5 and DT-K6) (Fig. 5c and Extended Data Fig. 5d), consistent with partial transcriptomic reprogramming toward an ATRT-SHH-like identity. Together, these findings demonstrate that *RFX4* cooperates with *SMARCB1* to adapt ATRT-MYC toward an ATRT-SHH transcriptomic state.

**Figure 5:**
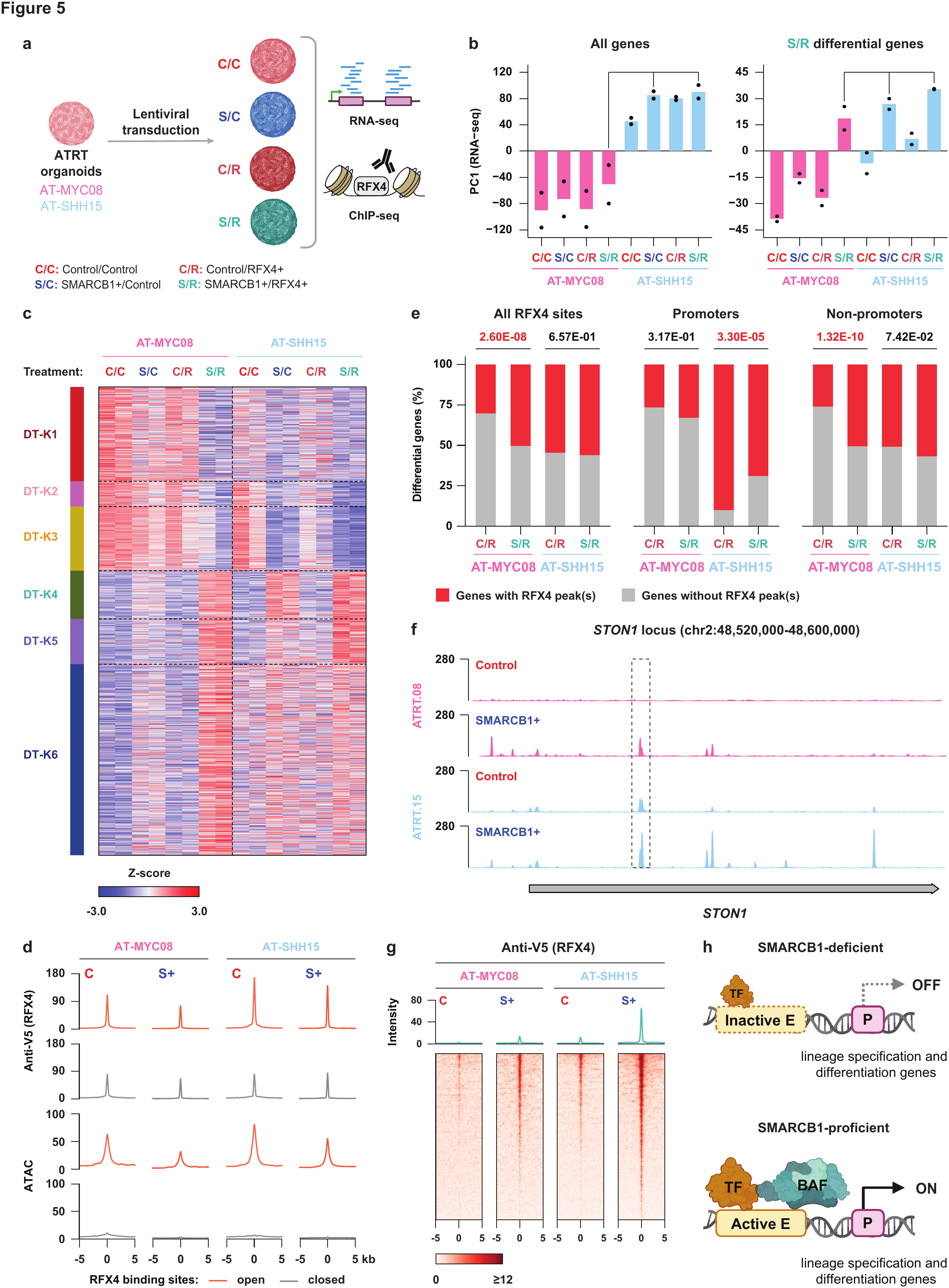
RFX4 and SMARCB1 cooperate to activate lineage-specific chromatin and transcription. a, Schematic overview of the dual lentiviral transduction strategy. C/C, Control/Control; S/C, SMARCB1+/Control; C/R, Control/RFX4+; S/R, SMARCB1+/RFX4+. **b,** Principal component analysis (PC1) of transcriptomic profiles from the dual transduction experiments using either all expressed genes (left) or differentially expressed genes identified in the AT-MYC08 S/R condition (right). **c,** Heatmap showing the six major expression patterns (DT-K1-DT-K6) encompassing 1,252 of the 1,328 differentially expressed genes identified in the AT-MYC08 S/R condition. **d,** Average RFX4 ChIP-seq (top two rows) and ATAC-seq (bottom two rows) signal centered on RFX4 binding sites (±5 kb) in AT-MYC08 and AT-SHH15 organoids without (C) or with (S+) *SMARCB1* reconstitution. Binding sites are stratified by chromatin accessibility (open, orange; closed, grey). **e,** Percentage of the differentially expressed genes identified in Extended Data Fig. 6a overlapping with all identified RFX4 binding sites, promoter associated RFX4 binding sites or RFX4 binding sites at non-promoter regulatory regions. P-values were calculated using Fisher’s exact test. **f,** An example of non-promoter RFX4 binding sites showing SMARCB1-dependent chromatin association in AT-MYC08. **g,** Average RFX4 ChIP-seq signal at K4 (SHH-up) differentially accessible regions identified by ATAC-seq (Fig. 3b). **h,** Proposed model illustrating how lineage-specific transcription factors cooperate with SMARCB1-containing BAF complexes to establish subtype-specific regulatory programs. In the context of ATRT (i.e., in the absence of SMARCB1), lineage-specific transcription factors retain limited occupancy at regulatory elements but fail to activate them because of impaired BAF complex function. Reconstitution of *SMARCB1* restores BAF activity, enabling transcription factor-directed chromatin remodeling and lineage-specific gene expression. Thus, the transcription factor repertoire specifies where BAF complexes act, whereas BAF complex integrity determines their regulatory output. E, enhancer; P, promoter.

To investigate how SMARCB1 influences RFX4 function at the chromatin level, we mapped RFX4 occupancy by ChIP-seq (Fig. 5a). Across both ATRT subtypes, *SMARCB1* reconstitution markedly increased the number of RFX4 binding sites compared with RFX4 expression alone (Extended Data Fig. 5e), demonstrating that intact BAF complexes facilitate RFX4 recruitment to chromatin. Moreover, the overlap in RFX4 binding sites between ATRT-MYC and ATRT-SHH increased substantially following *SMARCB1* reconstitution (Extended Data Fig. 5e), suggesting that restoration of BAF complex activity promotes a shared core RFX4 binding landscape. To determine whether RFX4 preferentially binds accessible chromatin, we stratified RFX4 binding sites according to their chromatin accessibility prior to *RFX4* expression. Surprisingly, RFX4 occupied both open and closed chromatin (Extended Data Fig. 5f), although binding at inaccessible regions was generally weaker than at accessible regions (Fig. 5d). This difference likely reflects the preferential localization of open RFX4 binding sites at promoter-proximal regions, whereas closed RFX4 binding sites were predominantly associated with distal regulatory elements (Extended Data Fig. 5g). Together, these findings suggest that RFX4 may possess pioneer chromatin-binding activity that enables its binding to both open and closed chromatin and further recruit SMARCB1 and BAF complex to ATRT subtype-specific regulatory elements. Next, we investigated the association between RFX4 chromatin occupancy and gene expression in AT-MYC08 and AT-SHH15 to understand whether chromatin regulatory functions of RXF4 contribute to transcriptomic adaption of ATRT-MYC toward ATRT-SHH in the SMARCB1-proficient condition. Because endogenous *RFX4* is exclusively expressed in ATRT-SHH (Extended Data Fig. 3f), we could expect that genes upregulated after *SMARCB1* and *RFX4* expression (S/R) have increased RFX4 binding sites in AT-MYC08 but not AT-SHH15. Therefore, we identified differentially expressed genes between the control (C/R) and SMARCB1+ (S/R) condition in the presence of RFX4 (Extended Data Fig. 6a), and interrogated enrichment of RFX4 binding sites associated with the differential genes in the Control (C/R) and the SMARCB1+ (S/R) conditions. As expected, the genes with higher expression in the S/R condition of AT-MYC08 also have increased percent of RFX4 binding sites especially those located at non-promoter regulatory regions (Fig. 5e), exemplified by a RFX4 binding sites in the intragenic region of *STON1* (Fig. 5f). In contrast, ectopic expression of RFX4 in AT-SHH15 did not increase the binding sites of RFX4 associated with S/R-specific genes (Fig. 5e).

Finally, we examined RFX4 occupancy at the six groups of SMARCB1-regulated differentially accessible regions (DARs) identified by ATAC-seq (Fig. 3b). As expected, *SMARCB1* reconstitution selectively increased chromatin accessibility at the SHH-specific K4 DARs in ATRT-SHH (Fig.3b and Extended Data Fig. 6b). Consistent with this increase in accessibility, RFX4 occupancy was strongly enriched at K4 DARs following *SMARCB1* reconstitution (Fig. 5g), exemplified by enhanced binding at putative enhancers downstream of *WNT7A* (Fig. 3e and Extended Data Fig. 6c). Notably, low-level RFX4 occupancy was already detectable at many K4 regulatory elements in the *SMARCB1*-deficient condition (Fig. 5g and Extended Data Fig. 6c), indicating that RFX4 can engage these lineage-specific sites before chromatin opening. In addition to K4, increased RFX4 binding was also observed at the commonly accessible K6 DARs in ATRT-SHH following *SMARCB1* reconstitution (Extended Data Fig. 6d), despite comparable chromatin accessibility at these regions in ATRT-MYC (Extended Data Fig. 6e). In contrast, RFX4 showed little or no binding across the remaining DAR classes (K1-K3 and K5), irrespective of their chromatin accessibility (Extended Data Fig. 6d).

Together, these findings support a model in which RFX4 and SMARCB1 cooperate to regulate lineage-specific regulatory elements, which may contribute to the unique molecular features and phenotypical traits of the ATRT-SHH subtype. Lineage-associated RFX4 binding at its target loci is weakly retained in the SMARCB1-deficient state, which may enable rapid activation of this regulatory elements upon BAF complex restoration and consequently activate transcription of lineage-specific genes (Fig. 5h). More broadly, these data demonstrate that the lineage-specific functions of the BAF complex are dictated by the underlying TF repertoire, providing a mechanistic explanation for how disruption of a single BAF complex subunit can give rise to distinct tumor identities.

## Discussion

BAF complexes are master regulators of chromatin accessibility and gene transcription, with highly context-dependent functions across developmental and cellular lineages. Genes encoding BAF complex subunits are frequently mutated in human cancer^54,55^, yet seemingly similar mutations can produce divergent regulatory outputs depending on cellular context. How disruption of BAF activity is translated into lineage-specific transcriptional programs that drive context-dependent malignant transformation remains a central unresolved question. Using ATRT as a genetically tractable model system, we show that *SMARCB1* inactivation does not define a single transcriptional state but instead enables divergent, lineage-restricted regulatory programs governed by the pre-existing TF repertoire of the cell.

We find that *SMARCB1* reconstitution restores a common BAF-dependent chromatin accessibility landscape while preserving subtype-specific transcriptomic identities. SMARCB1 therefore does not impose a uniform transcriptomic state but instead permits the existing TF repertoire to activate lineage-restricted regulatory programs in a similar cellular context. These results support a model in which BAF complexes act as context-dependent facilitators of developmental gene regulation, with tumor transcriptomic heterogeneity arising primarily from cellular state rather than the initiating genetic lesion.

Consistent with this model, we identify RFX4 as a lineage-restricted TF that is selectively engaged in ATRT-SHH regulatory programs upon restoration of SMARCB1. Single-nucleus multiome analyses of primary tumors reveal RFX4 as a key regulator of neuronal differentiation programs, with eRegulon activity enriched in ATRT-SHH and associated with more differentiated cell states^52^. Notably, the presence of RFX4 regulatory activity in ATRT-SHH tumors suggests that RFX4 retains at least some regulatory functions independent of SMARCB1. However, restoration of lineage-restricted transcriptomic programs requires cooperation between RFX4 and SMARCB1.

Functionally, ectopic expression of *RFX4* suppresses ATRT organoid growth regardless of subtype, likely due to its conserved functions in regulating cell cycle genes in neuronal lineages^49,56^. In the presence of SMARCB1, RFX4 activates lineage-associated gene programs and partially reprograms ATRT-MYC organoids toward an ATRT-SHH-like state. However, complete subtype conversion likely requires coordinated activity of multiple lineage transcription factors, including SOX2 and KLF12, rather than RFX4 alone. Together with prior studies showing that *SMARCB1* reconstitution enables enhancer engagement by lineage regulators such as TEAD1 and ZIC2 in ATRT-MYC^39^, these data suggest that *SMARCB1* loss broadly impairs the chromatin-dependent output of lineage-specific transcription factors, while the affected regulatory programs are determined by the context cells were in when *SMARCB1* was lost.

RFX4 exhibits properties consistent with a pioneer transcription factor, defined by the ability to engage nucleosomal DNA and initiate chromatin remodeling, as exemplified by GATA, OCT4, and SOX2 factors^57–60^. We find that RFX4 binds both accessible and inaccessible chromatin and exhibits weak occupancy at SHH-specific regulatory elements even in the absence of SMARCB1, indicating pre-established enhancer priming. Restoration of SMARCB1 markedly increases RFX4 occupancy and chromatin accessibility at these sites, enabling robust activation of the ATRT-SHH transcriptional program (Fig. 5h). SMARCB1 loss therefore disrupts BAF complex integrity and prevents full chromatin remodeling at primed regulatory elements, thereby locking cells in a poorly differentiated state. This model is consistent with recent work identifying the SWIFT (SWI/SNF Ig-fold for TF interactions) domain of SMARCD subunits as a conserved interface for lineage-specific TFs across cellular contexts^61^, providing a potential mechanistic basis for context-dependent recruitment of BAF complexes by factors such as RFX4.

In addition to subtype-specific regulatory programs, *SMARCB1* reconstitution induces subtype-independent changes in chromatin architecture, including reduced accessibility at CTCF motifs and increased enrichment of AP-1 motifs. This is consistent with preferential association of AP-1 with cBAF and CTCF with ncBAF complexes observed in, amongst others, ecMRTs^24,62–65^, suggesting that *SMARCB1* reconstitution rebalances BAF complex usage toward canonical cBAF-driven chromatin remodeling. These changes further align with reports that loss of CTCF occupancy accompanies activation of developmental gene programs during differentiation^66,67^.

Together, our results identify two determinants of ATRT molecular subtype identity: BAF complex integrity and the pre-existing TF repertoire in the cells undergoing malignant transformation. More broadly, this work serves as a mechanistic framework in which BAF complexes function as context-dependent chromatin regulators whose output is specified by lineage-restricted transcription factors. This principle may extend to other BAF-mutant malignancies, including synovial sarcoma^68,69^, lung cancer^70,71^ and leukemia^72,73^, where cellular context is likely to dictate the phenotypic consequences of BAF complex dysfunction.

## Materials and Methods

### Ethical statement

Written informed consent was provided by all patients and/or parents/guardians. ATRT samples were received under ethical approval of the ethics committee. Approval for Máxima patient samples and clinical data within the context of this study has been granted by the Máxima biobank and data access committee https://research.prinsesmaximacentrum.nl/en/core-facilities/biobank-clinical-research-committee (biobank nr. PMCLAB2018.005).

### Organoid culturing

ATRT organoids were cultured in tumor stem medium (TSM) in suspension plates as previously described^27^. Cells were passaged approximately every 7 days, and medium was refreshed every 3 - 4 days.

### Plasmid construction

pLKO.1-UbC-luciferase-blast and pLKO.1-UbC-hSMARCB1-blast lentiviruses were previously described^22,74^. RFX4 lentiviral expression plasmids (pCDH-EF1a-mScarlet-PGK-Puro, pCDH-EF1a-RFX4-PGK-Puro and pCDH-EF1a-RFX4(ΔDBD)-PGK-Puro) were generated by Gibson assembly using pCDH-EF1a-PGK-Puro (addgene #72484) as the backbone.

### Lentiviral transductions

Lentiviral transductions with pLKO.1-UbC-luciferase-blast^74^, pLKO.1-UbC-hSMARCB1-blast^22^, pCDH-EF1a-RFX4-PGK-Puro, pCDH-EF1a-RFX4(ΔDBD)-PGK-Puro or pCDH-EF1a-mScarlet-PGK-Puro lentiviruses were performed with a multiplicity of infection (MOI) of ± 0.4 as previously described^22,75,76^. After 48 hours, successfully transduced cells were selected using blasticidin (10 μg/ml) or puromycin (1 μg/mL).

### Immunohistochemistry

Organoids were fixed in 4% paraformaldehyde, dehydrated and embedded in paraffin. Sections were cut at a thickness of 4 μm and subjected to H&E or immunohistochemical anti-SMARCB1 staining (Cell Marque, 272-M18, predilute). For attached cells, a plate staining was performed using coated coverslips with 0.1% gelatin. Coverslips were fixed in 4% paraformaldehyde, and H&E or immunohistochemical staining was performed with anti-SMARCB1 antibody (Abcam, EPR20189).

### Bulk RNA-sequencing and analysis

RNA was isolated using TRIzol RNA isolation reagent (Invitrogen) according to the manufacturer’s instructions. RNA-sequencing (RNA-seq) libraries were prepared using Novogene NGS RNA Library Prep Set (PT042). The libraries were sequenced on Novaseq 6000 platform with PE150 (Illumina). Two biological replicates were generated for each of the RNA-seq experiments in this study.

Pre-processing of raw reads was performed as previously described^24^. The recorded RNA-seq data were mapped to human reference genome (hg38) using TopHat2 with “--coverage-search” mode (version 2.1.1)^77^. The reads with mapping quality of at least 10 were kept for the subsequent analysis. Gene expression was determined by counting read coverage of the hg38 genes using HTSeq with non-stranded and union mode (version 1.99.2)^78^. The gene reads were normalized and then differentially expressed genes were detected using DESeq2 (version 1.49.4)^79^. To determine common and subtype specific transcriptomic changes, differentially expressed genes between the Control and SMARCB1+ condition were first determined within each of the two subtypes, and subsequently the genes affected either in both subtypes, or in one of the two subtypes were categorized into six K-mean clusters. For data visualization, Circos plots and heatmaps were plotted using circlize (version 0.4.16) and gplots (version 3.2.0), respectively. Other types of plots were visualized using ggplot2 (version 4.0.0). Gene ontology analysis was performed using g:Profiler^80^and Gene Set Enrichment Analysis was performed using clusterProfiler (version 4.16.0)^81^ and msigdbr (version 25.1.1)^82^.

### Bulk assay for transposase-accessible chromatin (ATAC)-sequencing and analysis

Organoids were harvested, washed with PBS and dead cells were removed using Dead Cell Removal Kit (Miltenyi Biotec). Library preparation was performed as previously described^24^. In short, nuclei were isolated and permeabilized. The isolated nuclei were tagmented using Tn5 transposase (Illumina, Nextera DNA Library Preparation Kit), followed by two sequential 9-cycle PCR. The resulting DNA fragments (<700 bp) were purified using SPRI beads (Beckman). ATAC-seq libraries were sequenced on Novaseq 6000 platform with PE150 (Illumina). All ATAC-seq experiments were performed in duplicates.

ATAC-seq reads were mapped to human reference genome (hg38) with BWA-MEM algorithm using bwa (version 0.7.17-r1188)^83^. The reads with mapping quality of at least 10 were kept and optical PCR duplicated reads were removed. Open chromatin sites, or peaks, were detected using macs2 (version 2.1.4)^84^ with narrow mode and at a q value cutoff of 0.01. For visualization, the bigWig files of the ATAC-seq data were generated using deeptools (version 3.5.5)^85^. Chromatin accessibility was determined by counting read coverage of the identified ATAC-seq peaks using HTSeq with union mode (version 1.99.2)^78^. Data normalization, detection of differentially accessible regions and data visualization were performed using the same R packages as used in RNA-seq analysis, except for visualizing common and subtype specific open chromatin changes which was performed using heatmap3 (version 1.1.9)^86^. Functional annotation of the differential open chromatin sites was performed using ChIPseeker (version 1.46.1)^87^. ATAC-seq footprinting was performed using TOBIAS (version 0.17.0)^88^.

### CellTiter-Glo assays

CellTiter-Glo (CTG) assays were performed according to manufacturer’s instructions. Cell growth was assessed at T0 (day of plating after selection of transduced cells) and either five (T5) or seven (T7) after plating.

### Western Blot

Cells were harvested and lysed in RIPA buffer (50 mM Tris-HCl pH 8.0, 150 mM NaCl, 0.1% SDS, 0.5% Na-Deoxycholate, 1% NP-40) followed by sonication (30 seconds on, 30 seconds off, 10 cycles in total) of the lysate using Bioruptor® Pico. Western blot was performed as previously described^89^. Nitrocellulose membranes (Whatman^TM^, #10600015) were blocked in 5% BSA in TBS-Tween (0.1%) and probed with primary antibodies against SMARCB1 (A-5, Santa Cruz Biotechnologies, #G1916), V5 (Life technologies, #R960-25), RFX4 (ThermoFisher, #PA5-62205) and GAPDH (Abcam, #ab-9485).

### snMultiome data processing and analysis

Data generation and preprocessing were performed as previously described^24,52^, with minor modifications as described below. ArchR (version 1.0.3)^90^ was used for data analysis. Cells were filtered to retain good quality (ATAC counts <100000 & > 1000, RNA counts <25000 & > 1000, nucleosome signal <2, TSS enrichment > 1). In ArchR, log-normalization was used for the snRNA-seq data and term frequency-inverse document frequency (TF-IDF) normalization was used for snATAC-seq data. Dimensionality reduction was performed using an Iterative LSI approach. Uniform Manifold Approximation Projection (UMAP) embeddings were generated using combined reduced dimensions (ArchR default, dimensions 1:60). Motif deviations were calculated per cell using the ArchR implementation of ChromVAR (version 1.28.0)^91^ and CIS-BP derived motif collection curated by the chromVARmotifs package (ArchR default, version 2). Gene regulatory networks were inferred based on the snMultiome data using SCENIC+ (version 1.0a2)^92^. For the SCENIC+ pipeline, data was pre-processed using pycisTopic (version 2.0a0) and pycistarget (version 1.1). The pycisTopic model with 70 topics was selected for further processing. SCENIC+ output was filtered for the top 5 ‘+/+’ eRegulons per ATRT sample, which refer to a positive correlation between TF and the accessibility of its regulatory regions and gene expression of its target genes. Gene-based and region-based Area Under the Curve (AUC) scores were extracted and averaged per sample and further scaled across all samples for visualization.

### Double crosslinked chromatin immunoprecipitation (ChIP)-sequencing

The ChIP-seq experiments were performed as previously described^93^. Double crosslinking was carried out in two sequential steps. Organoids were first harvested, washed in PBS and treated with 2 mM disuccinimidyl glutarate (DSG, 20593, Thermo Fisher Scientific) for 45 minutes at room temperature, followed by a single wash with PBS. A second crosslinking step was then performed by exposure to 37% formaldehyde for 10 minutes. Excess crosslinker was quenched by addition of 2.0 M glycine. Crosslinked cells were subsequently lysed, and chromatin was sheared by sonication for 5 minutes with a “30s-on-30s-off” mode using a Bioruptor Plus device (Diagenode). For ChIP, antibodies were pre-coupled to Protein G magnetic beads (Thermo Fisher Scientific) prior to overnight incubation with sheared chromatin at 4°C. Immunoprecipitated chromatin complexes were sequentially washed, eluted, and subjected to reverse crosslinking. DNA was recovered from the eluate using the MiniElute PCR Purification Kit (Qiagen). The following antibodies were used: V5 (Life technologies, #R960-25, 5 μl per ChIP). The purified DNA fragments were prepared using the KAPA HTP Library Preparation Kit (Roche) following manufacturer’s instructions. The libraries were sequenced on an Illumina NovaSeq 6000 platform using the paired-end 100-cycle mode.

ChIP-seq data were analyzed as previously described^24^. In short, sequencing reads were mapped to the hg38 reference genome using the Bowtie 2 mapper (version 2.3.4.1)^94^. The mapped reads were filtered using SAMtools, discarding reads with mapping quality score <15, as well as optical PCR duplicates. The coverage files were produced using the deepTools (version 3.0) method “normalize to 1X genome coverage”^85^. Functional annotation of the differential open chromatin sites was performed using ChIPseeker (version 1.46.1)^87^. Assignment of ChIP-seq peaks to their gene targets was performed using the “basal plus extension” method^95^ in the rGREAT package (version 2.12.1)^96^. ChIP-seq peaks overlap with transcription start sites extending ± 1 kb of annotated genes were considered as promoter binding sites of RFX4, and the rest of the peaks were defined as non-promoter binding sites.

## Data availability statements

Next-generation sequencing data that support the findings of this study have been deposited in Gene Expression Omnibus and are accessible through GEO Series accession number GSEXXXXXX.

## Acknowledgments

We are deeply grateful to the patients and their families for participating in this study, without whose support this work would not have been possible. We thank the Single-Cell Genomics Facility at the Princess Máxima Center, the Genomic Core Facility in Department of Hematology at the Erasmus Medical Center Cancer Institute and the Genomics Core Facility at the NKI, as well as the contributions of the Princess Máxima Center Biobank and Data Access Committee.

## Author Contributions

J.D., N.Q.L. and E.d.W. supervised the work. J.D., N.Q.L. and M.K. contributed to funding resources. J.H. and C.R.A. performed the experimental work with technical support from N.A., W.T. and I.P. N.Q.L. and S.J. performed data analyses. J.H., C.R.A., N.Q.L. and J.D. wrote the manuscript with input from all co-authors.

## References

1. Shain, A. H. & Pollack, J. R. The spectrum of SWI/SNF mutations, ubiquitous in human cancers. PLoS One 8, e55119 (2013).

2. Kadoch, C. et al. Proteomic and bioinformatic analysis of mammalian SWI/SNF complexes identifies extensive roles in human malignancy. Nat. Genet. 45, 592–601 (2013).

3. Church, M. C. & Workman, J. L. The SWI/SNF chromatin remodeling complex: a critical regulator of metabolism. Biochem. Soc. Trans. 52, 1327–1337 (2024).

4. Centore, R. C., Sandoval, G. J., Soares, L. M. M., Kadoch, C. & Chan, H. M. Mammalian SWI/SNF Chromatin Remodeling Complexes: Emerging Mechanisms and Therapeutic Strategies. Trends Genet. 36, 936–950 (2020).

5. Kadoch, C. & Crabtree, G. R. Mammalian SWI/SNF chromatin remodeling complexes and cancer: Mechanistic insights gained from human genomics. Sci. Adv. 1, e1500447 (2015).

6. Varga, J., Kube, M., Luck, K. & Schick, S. The BAF chromatin remodeling complexes: structure, function, and synthetic lethalities. Biochem. Soc. Trans. 49, 1489–1503 (2021).

7. Hota, S. K. et al. Dynamic BAF chromatin remodeling complex subunit inclusion promotes temporally distinct gene expression programs in cardiogenesis. Development 146, (2019).

8. Oo, J. A. et al. Long non-coding RNAs direct the SWI/SNF complex to cell type-specific enhancers. Nat. Commun. 16, 131 (2025).

9. Lara-Astiaso, D. et al. In vivo screening characterizes chromatin factor functions during normal and malignant hematopoiesis. Nat. Genet. 55, 1542–1554 (2023).

10. Marei, H. E. Epigenetic regulators in cancer therapy and progression. *NPJ Precis*. Oncol. 9, 206 (2025).

11. Gröbner, S. N. et al. The landscape of genomic alterations across childhood cancers. Nature 555, 321–327 (2018).

12. Ma, X. et al. Pan-cancer genome and transcriptome analyses of 1,699 paediatric leukaemias and solid tumours. Nature 555, 371–376 (2018).

13. Lee, R. S. et al. A remarkably simple genome underlies highly malignant pediatric rhabdoid cancers. J. Clin. Invest. 122, 2983–8 (2012).

14. Chun, H.-J. E. et al. Identification and Analyses of Extra-Cranial and Cranial Rhabdoid Tumor Molecular Subgroups Reveal Tumors with Cytotoxic T Cell Infiltration. Cell Rep. 29, 2338–2354.e7 (2019).

15. Ho, B. et al. Molecular subgrouping of atypical teratoid/rhabdoid tumors-a reinvestigation and current consensus. Neuro. Oncol. 22, 613–624 (2020).

16. Hasselblatt, M. et al. SMARCA4-mutated atypical teratoid/rhabdoid tumors are associated with inherited germline alterations and poor prognosis. Acta Neuropathol. 128, 453–6 (2014).

17. Tang, M. & Verhaak, R. G. A Molecular Take on Malignant Rhabdoid Tumors. Trends Cancer 2, 217–218 (2016).

18. Nemes, K. et al. Infants and Newborns with Atypical Teratoid Rhabdoid Tumors (ATRT) and Extracranial Malignant Rhabdoid Tumors (eMRT) in the EU-RHAB Registry: A Unique and Challenging Population. Cancers (Basel). 14, (2022).

19. Johann, P. D. et al. Atypical Teratoid/Rhabdoid Tumors Are Comprised of Three Epigenetic Subgroups with Distinct Enhancer Landscapes. Cancer Cell 29, 379–393 (2016).

20. Han, Z. Y. et al. The occurrence of intracranial rhabdoid tumours in mice depends on temporal control of Smarcb1 inactivation. Nat. Commun. 7, (2016).

21. Roberts, C. W. M., Leroux, M. M., Fleming, M. D. & Orkin, S. H. Highly penetrant, rapid tumorigenesis through conditional inversion of the tumor suppressor gene Snf5. Cancer Cell 2, 415–25 (2002).

22. Custers, L. et al. Somatic mutations and single-cell transcriptomes reveal the root of malignant rhabdoid tumours. Nat. Commun. 12, 1407 (2021).

23. Vitte, J., Gao, F., Coppola, G., Judkins, A. R. & Giovannini, M. Timing of Smarcb1 and Nf2 inactivation determines schwannoma versus rhabdoid tumor development. Nat. Commun. 8, 300 (2017).

24. Liu, N. Q. et al. SMARCB1 loss activates patient-specific distal oncogenic enhancers in malignant rhabdoid tumors. Nat. Commun. 14, 7762 (2023).

25. Wang, X. et al. SMARCB1-mediated SWI/SNF complex function is essential for enhancer regulation. Nat. Genet. 49, 289–295 (2017).

26. Nakayama, R. T. et al. SMARCB1 is required for widespread BAF complex-mediated activation of enhancers and bivalent promoters. Nat. Genet. 49, 1613–1623 (2017).

27. Paassen, I. et al. Atypical teratoid/rhabdoid tumoroids reveal subgroup-specific drug vulnerabilities. Oncogene 42, 1661–1671 (2023).

28. Nakayama, R. T. et al. SMARCB1 is required for widespread BAF complex-mediated activation of enhancers and bivalent promoters. Nat. Genet. 49, 1613–1623 (2017).

29. Torchia, J. et al. Integrated (epi)-Genomic Analyses Identify Subgroup-Specific Therapeutic Targets in CNS Rhabdoid Tumors. Cancer Cell 30, 891–908 (2016).

30. Centore, R. C., Sandoval, G. J., Soares, L. M. M., Kadoch, C. & Chan, H. M. Mammalian SWI/SNF Chromatin Remodeling Complexes: Emerging Mechanisms and Therapeutic Strategies. Trends Genet. 36, 936–950 (2020).

31. Borsari, B. et al. Enhancers with tissue-specific activity are enriched in intronic regions. Genome Res. 31, 1325–1336 (2021).

32. Panigrahi, A. & O’Malley, B. W. Mechanisms of enhancer action: the known and the unknown. Genome Biol. 22, 108 (2021).

33. Nakayama, R. T. et al. SMARCB1 is required for widespread BAF complex-mediated activation of enhancers and bivalent promoters. Nat. Genet. 49, 1613–1623 (2017).

34. Wang, X. et al. SMARCB1-mediated SWI/SNF complex function is essential for enhancer regulation. Nat. Genet. 49, 289–295 (2017).

35. Erkek, S. et al. Comprehensive Analysis of Chromatin States in Atypical Teratoid/Rhabdoid Tumor Identifies Diverging Roles for SWI/SNF and Polycomb in Gene Regulation. Cancer Cell 35, 95–110.e8 (2019).

36. Huhtala, L., Karabiyik, G. & Rautajoki, K. J. Development and epigenetic regulation of Atypical teratoid/rhabdoid tumors in the context of cell-of-origin and halted cell differentiation. Neurooncol. Adv. 6, vdae162 (2024).

37. Graf, M. et al. Single-cell transcriptomics identifies potential cells of origin of MYC rhabdoid tumors. Nat. Commun. 13, 1544 (2022).

38. Kenny, C., O’Meara, E., Ulaş, M., Hokamp, K. & O’Sullivan, M. J. Global Chromatin Changes Resulting from Single-Gene Inactivation-The Role of SMARCB1 in Malignant Rhabdoid Tumor. Cancers (Basel). 13, (2021).

39. Nesvick, C. L. et al. SWI/SNF complexes govern ontology-specific transcription factor function in MYC-subtype atypical teratoid rhabdoid tumor. Neuro. Oncol. 27, 2445–2460 (2025).

40. Hettige, N. C. & Ernst, C. FOXG1 Dose in Brain Development. Front. Pediatr. 7, 482 (2019).

41. Hubert, K. A. & Wellik, D. M. Hox genes in development and beyond. Development 150, (2023).

42. Co, M., Anderson, A. G. & Konopka, G. FOXP transcription factors in vertebrate brain development, function, and disorders. Wiley Interdiscip. Rev. Dev. Biol. 9, e375 (2020).

43. Hettige, N. C. & Ernst, C. FOXG1 Dose in Brain Development. Front. Pediatr. 7, 482 (2019).

44. Rousso, D. L. et al. Foxp-mediated suppression of N-cadherin regulates neuroepithelial character and progenitor maintenance in the CNS. Neuron 74, 314–30 (2012).

45. Philippidou, P. & Dasen, J. S. Hox genes: choreographers in neural development, architects of circuit organization. Neuron 80, 12–34 (2013).

46. Charité, J. et al. Transducing positional information to the Hox genes: critical interaction of cdx gene products with position-sensitive regulatory elements. Development 125, 4349–58 (1998).

47. Bergsland, M. et al. Sequentially acting Sox transcription factors in neural lineage development. Genes Dev. 25, 2453–64 (2011).

48. Stevanovic, M. et al. SOX Transcription Factors as Important Regulators of Neuronal and Glial Differentiation During Nervous System Development and Adult Neurogenesis. Front. Mol. Neurosci. 14, 654031 (2021).

49. Choi, W. et al. RFX4 is an intrinsic factor for neuronal differentiation through induction of proneural genes POU3F2 and NEUROD1. Cell. Mol. Life Sci. 81, (2024).

50. Sugiaman-Trapman, D. et al. Characterization of the human RFX transcription factor family by regulatory and target gene analysis. BMC Genomics 19, 1–15 (2018).

51. Ashique, A. M. et al. The Rfx4 transcription factor modulates Shh signaling by regional control of ciliogenesis. Sci. Signal. 2, ra70 (2009).

52. Blanco-Carmona, E. et al. A cycling, progenitor-like cell population at the base of atypical teratoid rhabdoid tumor subtype differentiation trajectories. Neuro. Oncol. (2025) doi:10.1093/neuonc/noaf179.

53. Smith, H. L., Aouad, P. & Wadhwani, N. R. Atypical Teratoid Rhabdoid Tumor: How Tumor Diagnostic Methods in the Laboratory Have Evolved over the Past 40 Years. Cancers (Basel). 17, (2025).

54. Kadoch, C. et al. Proteomic and bioinformatic analysis of mammalian SWI/SNF complexes identifies extensive roles in human malignancy. Nat. Genet. 45, 592–601 (2013).

55. Shain, A. H. & Pollack, J. R. The spectrum of SWI/SNF mutations, ubiquitous in human cancers. PLoS One 8, e55119 (2013).

56. Determan, J. J. et al. Intellectual disability risk gene RFX4 regulates cortical neurogenesis by restraining neuronal differentiation. Preprint at 10.64898/2026.02.11.705165 (2026).

57. Mayran, A. & Drouin, J. Pioneer transcription factors shape the epigenetic landscape. J. Biol. Chem. 293, 13795–13804 (2018).

58. Soufi, A. et al. Pioneer transcription factors target partial DNA motifs on nucleosomes to initiate reprogramming. Cell 161, 555–568 (2015).

59. Iwafuchi-Doi, M. The mechanistic basis for chromatin regulation by pioneer transcription factors. Wiley Interdiscip. Rev. Syst. Biol. Med. 11, e1427 (2019).

60. Zaret, K. S. & Carroll, J. S. Pioneer transcription factors: establishing competence for gene expression. Genes Dev. 25, 2227–41 (2011).

61. Jain, S. U. et al. A SWI/SNF-specific Ig-like domain, SWIFT, is a transcription factor binding platform. Science 392, eaeb3627 (2026).

62. Michel, B. C. et al. A non-canonical SWI/SNF complex is a synthetic lethal target in cancers driven by BAF complex perturbation. Nat. Cell Biol. 20, 1410–1420 (2018).

63. Xiao, M. et al. BRD9 determines the cell fate of hematopoietic stem cells by regulating chromatin state. Nat. Commun. 14, 8372 (2023).

64. Vierbuchen, T. et al. AP-1 Transcription Factors and the BAF Complex Mediate Signal-Dependent Enhancer Selection. Mol. Cell 68, 1067–1082.e12 (2017).

65. Otto, J. E. et al. Structural and functional properties of mSWI/SNF chromatin remodeling complexes revealed through single-cell perturbation screens. Mol. Cell 83, 1350–1367.e7 (2023).

66. Michel, B. C. et al. A non-canonical SWI/SNF complex is a synthetic lethal target in cancers driven by BAF complex perturbation. Nat. Cell Biol. 20, 1410–1420 (2018).

67. Liu, Y. et al. CTCF coordinates cell fate specification via orchestrating regulatory hubs with pioneer transcription factors. Cell Rep. 42, 113259 (2023).

68. Brien, G. L. et al. Targeted degradation of BRD9 reverses oncogenic gene expression in synovial sarcoma. Elife 7, (2018).

69. McBride, M. J. et al. The SS18-SSX Fusion Oncoprotein Hijacks BAF Complex Targeting and Function to Drive Synovial Sarcoma. Cancer Cell 33, 1128–1141.e7 (2018).

70. Duplaquet, L. et al. Mammalian SWI/SNF complex activity regulates POU2F3 and constitutes a targetable dependency in small cell lung cancer. Cancer Cell 42, 1352–1369.e13 (2024).

71. de Miguel, F. J. et al. Mammalian SWI/SNF chromatin remodeling complexes promote tyrosine kinase inhibitor resistance in EGFR-mutant lung cancer. Cancer Cell 41, 1516–1534.e9 (2023).

72. Del Gaudio, N. et al. BRD9 binds cell type-specific chromatin regions regulating leukemic cell survival via STAT5 inhibition. Cell Death Dis. 10, 338 (2019).

73. Hohmann, A. F. et al. Sensitivity and engineered resistance of myeloid leukemia cells to BRD9 inhibition. Nat. Chem. Biol. 12, 672–9 (2016).

74. Fumagalli, A. et al. Genetic dissection of colorectal cancer progression by orthotopic transplantation of engineered cancer organoids. Proc. Natl. Acad. Sci. U. S. A. 114, E2357–E2364 (2017).

75. Koo, B.-K. et al. Controlled gene expression in primary Lgr5 organoid cultures. Nat. Methods 9, 81–3 (2011).

76. Drost, J., Artegiani, B. & Clevers, H. The generation of organoids for studying Wnt signaling. Methods in Molecular Biology 1481, 141–159 (2016).

77. Kim, D. et al. TopHat2: accurate alignment of transcriptomes in the presence of insertions, deletions and gene fusions. Genome Biol. 14, (2013).

78. Putri, G. H., Anders, S., Pyl, P. T., Pimanda, J. E. & Zanini, F. Analysing high-throughput sequencing data in Python with HTSeq 2.0. Bioinformatics 38, 2943–2945 (2022).

79. Love, M. I., Huber, W. & Anders, S. Moderated estimation of fold change and dispersion for RNA-seq data with DESeq2. Genome Biol. 15, (2014).

80. Reimand, J., Kull, M., Peterson, H., Hansen, J. & Vilo, J. g:Profiler--a web-based toolset for functional profiling of gene lists from large-scale experiments. Nucleic Acids Res. 35, W193–200 (2007).

81. Yu, G., Wang, L.-G., Han, Y. & He, Q.-Y. clusterProfiler: an R package for comparing biological themes among gene clusters. OMICS 16, 284–7 (2012).

82. Liberzon, A. et al. The Molecular Signatures Database (MSigDB) hallmark gene set collection. Cell Syst. 1, 417–425 (2015).

83. Li, H. & Wren, J. Toward better understanding of artifacts in variant calling from high-coverage samples. Bioinformatics 30, 2843–2851 (2014).

84. Zhang, Y. et al. Model-based analysis of ChIP-Seq (MACS). Genome Biol. 9, R137 (2008).

85. Ramírez, F., Dündar, F., Diehl, S., Grüning, B. A. & Manke, T. deepTools: a flexible platform for exploring deep-sequencing data. Nucleic Acids Res. 42, (2014).

86. Zhao, S., Guo, Y., Sheng, Q. & Shyr, Y. Advanced heat map and clustering analysis using heatmap3. Biomed Res. Int. 2014, 986048 (2014).

87. Yu, G., Wang, L.-G. & He, Q.-Y. ChIPseeker: an R/Bioconductor package for ChIP peak annotation, comparison and visualization. Bioinformatics 31, 2382–3 (2015).

88. Bentsen, M. et al. ATAC-seq footprinting unravels kinetics of transcription factor binding during zygotic genome activation. Nat. Commun. 11, 4267 (2020).

89. Drost, J. et al. Sequential cancer mutations in cultured human intestinal stem cells. Nature 2015 521:7550 521, 43–47 (2015).

90. Granja, J. M. et al. ArchR is a scalable software package for integrative single-cell chromatin accessibility analysis. Nat. Genet. 53, 403–411 (2021).

91. Schep, A. N., Wu, B., Buenrostro, J. D. & Greenleaf, W. J. chromVAR: inferring transcription-factor-associated accessibility from single-cell epigenomic data. Nat. Methods 14, 975–978 (2017).

92. Bravo González-Blas, C., et al. SCENIC+: single-cell multiomic inference of enhancers and gene regulatory networks. Nat. Methods 20, 1355–1367 (2023).

93. Liu, N. Q. et al. WAPL maintains a cohesin loading cycle to preserve cell-type-specific distal gene regulation. Nat. Genet. 53, 100–109 (2021).

94. Langmead, B., Trapnell, C., Pop, M. & Salzberg, S. L. Ultrafast and memory-efficient alignment of short DNA sequences to the human genome. Genome Biol. 10, R25 (2009).

95. McLean, C. Y. et al. GREAT improves functional interpretation of cis-regulatory regions. Nat. Biotechnol. 28, 495–501 (2010).

96. Gu, Z. & Hübschmann, D. rGREAT: an R/bioconductor package for functional enrichment on genomic regions. Bioinformatics 39, (2023).

