## Supplementary Figures for "BAF complexes cooperate with lineage-restricted transcription factors to establish cell-state-specific regulatory programs"

### Extended Data

**Extended Data Figure 1: Phenotypic characterization of ATRT organoids upon *SMARCB1* reconstitution. a, b,** Representative images of INI-1 (**a**) and hematoxylin and eosin (H&E) (**b**) stainings on the indicated ATRT organoid models with Control and SMARCB1+ transduction. Scale bars = 100 µm. **c,** Representative brightfield images of the indicated Control and SMARCB1+ ATRT models. Scale bars = 100 µm.

**Extended Data Figure 2: Transcriptomic effects of *SMARCB1* reconstitution in the two ATRT subtypes. a,** Box plot of normalized reads of *SMARCB1* in all seven organoid models (3x SHH and 4x MYC) in the Control (C) and SMARCB1+ (S+) conditions. **b,** Principal component analysis (PCA) uncovering transcriptome-wide changes following *SMARCB1* reconstitution. **c,** Bar plot summarizing the differentially expressed genes induced by *SMARCB1* reconstitution in each of the isogenic organoid pairs. **d,** Boxplots of normalized gene expression (normalized reads, log2) per gene cluster (K1, K4, K5 or K6) in MYC and SHH subtypes. C = Control; S+ = SMARCB1+. **e,** A model summarizing how *SMARCB1* loss may contribute to subtype-specific transcriptomic landscape of ATRT**.** *SMARCB1* loss in the cell of origin does not alter the pre-existing transcription factor repertoire (Extended Data Fig. 2d) but instead prevents this repertoire from activating its associated lineage-specifying gene programs. In ATRT-MYC, this manifests as failure to induce SHH-associated genes (K4 cluster, e.g. *FGF1*), whereas in ATRT-SHH, MYC-associated genes (K5 cluster, e.g. *MYCN*) remain unexpressed. We propose that this asymmetric block, rather than an instructive transcriptional program, underlies the divergent identities of the two ATRT subtypes.

**Extended Data Figure 3: Open chromatin profiling identifies subtype-specific TF regulators cooperating with SMARCB1 in ATRT. a,** PCA uncovering genome-wide open chromatin landscape in the Control and SMARCB1+ organoids. **b,** Barplot summarizing the DARs induced by *SMARCB1* reconstitution in each of the six isogenic organoid pairs. **c,** Stacked bar plot showing the genomic distribution of open chromatin regions across the 6 open chromatin clusters. All open chromatin sites (All) serve as the background to check genome average distribution of accessible chromatin. **d,** Volcano plots of genome-wide TF footprinting analysis per ATRT organoid model, where every dot represents an individual TF motif. The x-axis represents differential binding score (Control versus SMARCB1+), and the y-axis indicates statistical significance. **e,** Tn5 digestion pattern around the RFX motifs (footprints) of the K4 (SHH up) open chromatin regions in the six ATRT organoid models. Aggregate binding signals are plotted ±60 bp from motif centers, with dashed lines indicating the edge of RFX motifs. **f,** Box plots showing the expression of all the expressed *RFX* genes in the isogenic organoid pairs. C = Control; S+ = SMARCB1+.

**Extended Data Figure 4: RFX4 is an ATRT-SHH-specific transcription factor that impairs growth. a,** UMAP projection of single-nucleus transcriptomic profiles colored by *RFX4* expression levels. Color intensity indicating normalized expression (normalized counts). **b,** UMAP projection of single-nucleus data colored by gene AUC scores for the identified RFX3 eRegulon. **c,** UMAP projection showing the cell distribution of the ATRT-SHH cells in the top 25% (marked red, named "RFX4-high") and bottom 25% (marked dark blue, named "RFX4-low") of RFX4 eRegulon activity. **d,** A PCA plot of transcriptome-wide changes following ectopic RFX4 expression in 2x ATRT-SHH and 2x ATRT-MYC models.

**Extended Data Figure 5: Exogenous *RFX4* expression and SMARCB1 reactivation are critical for maintaining lineage identity of ATRT-SHH. a,** PCA of transcriptomic changes in all genes (left) or differentially expressed genes identified in the AT-MYC08 S/R condition (right). C/C = control/control; S/C = SMARCB1+/Control; C/R = Control/RFX4+; S/R = SMARCB1+/RFX4+. **b,** Bar plot of differentially expressed genes in the RFX4 and/or SMARCB1 transduced conditions compared to the double Control condition (C/C, as reference in the statistical tests). **c,** Expression fold changes (compared with C/C, horizontal dashed lines) of the 6 gene clusters (Fig. 2c) in the dual transduction experiments. **d,** Gene Ontology analysis of the six DT-K gene clusters showing representative enriched biological processes. Enrichments are classified as common (present in ≥2 clusters) or cluster-specific (DT-K1, DT-K2, DT-K3, DT-K5, and DT-K6). Scale bar indicates statistical significance (−log10 FDR). **e,** Venn diagrams showing unique and shared RFX4 binding sites between AT-MYC08 and AT-SHH15 organoids under Control (top) and SMARCB1+ (bottom) conditions. Pale violet red indicates AT-MYC08-specific peaks, sky blue indicates AT-SHH15-specific peaks, and grey indicates shared peaks. **f,** Pie charts showing distribution of RFX4 binding sites under Control (C) and SMARCB1+ (S+) conditions, stratified by chromatin accessibility as determined by ATAC-seq. Binding sites are classified as occurring in closed (grey) or open (orange) chromatin. **g,** Stacked bar plot showing the genomic distribution of the RFX4-V5 ChIP-seq peaks from the Control and SMARCB1+ conditions of AT-MYC08 and AT-SHH15, stratified by open (O) or closed (C) chromatin as observed in ATAC-seq. All the ChIP-seq peaks serve as the background to check genome average distribution of RFX4-V5 binding sites.

**Extended Data Figure 6: Exogenous *RFX4* expression and SMARCB1 reactivation are critical for maintaining lineage identity of ATRT-SHH. a,** Volcano plots showing differentially expressed genes between the Control (C/R) and SMARCB1+ (S/R) condition in the presence of exogenous RFX4. **b,** Quantification of chromatin accessibility at the K4 (SHH up) open chromatin sites from ATAC-seq (Fig. 3b). **c,** An example of RFX4-binding locus (*WNT7A*) exclusive to ATRT-SHH, increased upon SMARCB1 restoration. The dashed box highlights a regulatory element containing a RFX motif which gains chromatin accessibility upon SMARCB1 reactivation in ATRT-SHH (Fig. 3e). **d,** Average RFX4-V5 binding profile at the K1/K2/K3/K5/K6 open chromatin sites identified by ATAC-seq (Fig. 3b). **e,** Quantification of chromatin accessibility at the K6 (all up) open chromatin sites from ATAC-seq (Fig. 3b).

Extended Data Fig. 1

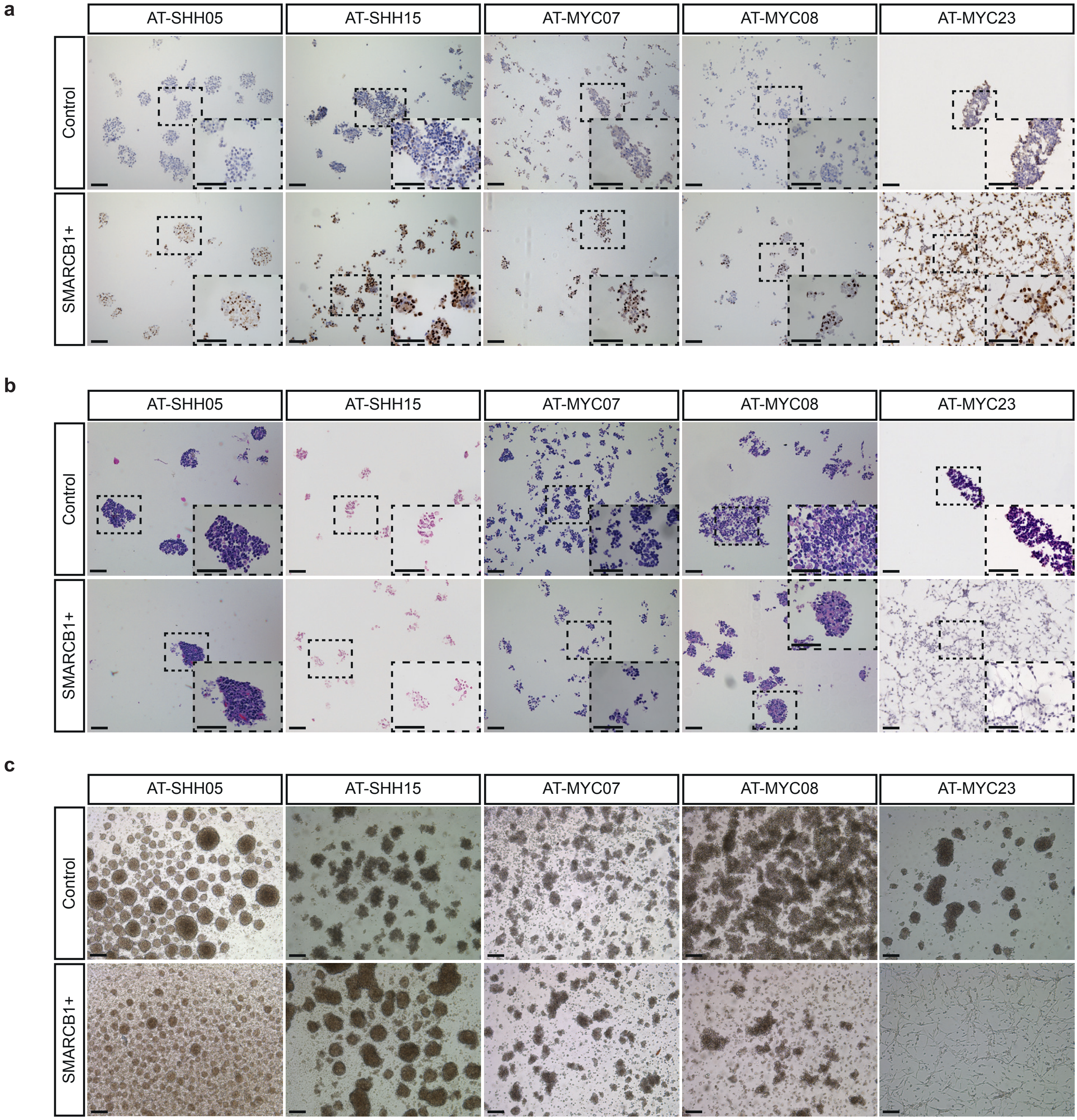

Extended Data Fig. 2

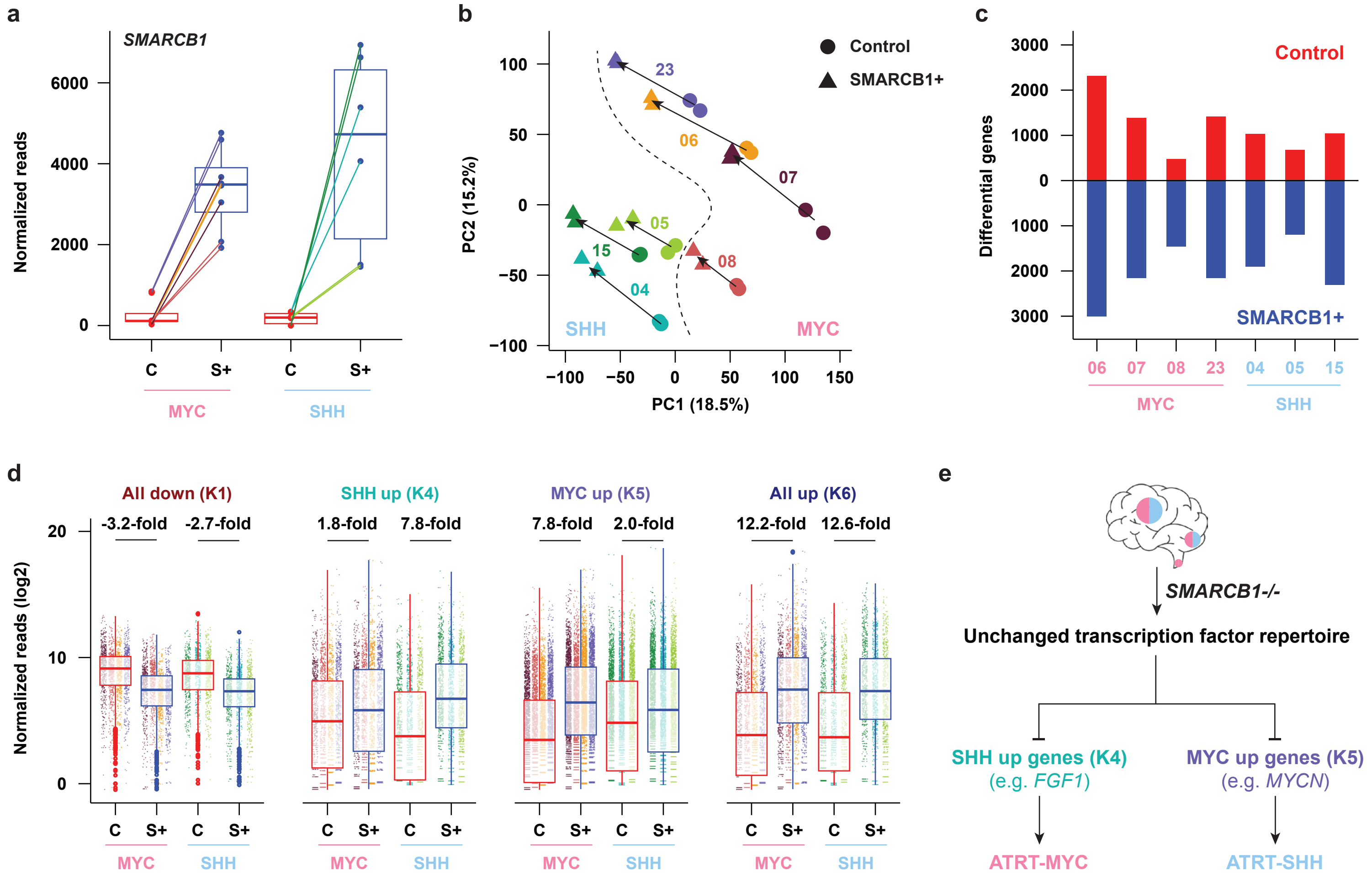

Extended Data Fig. 3

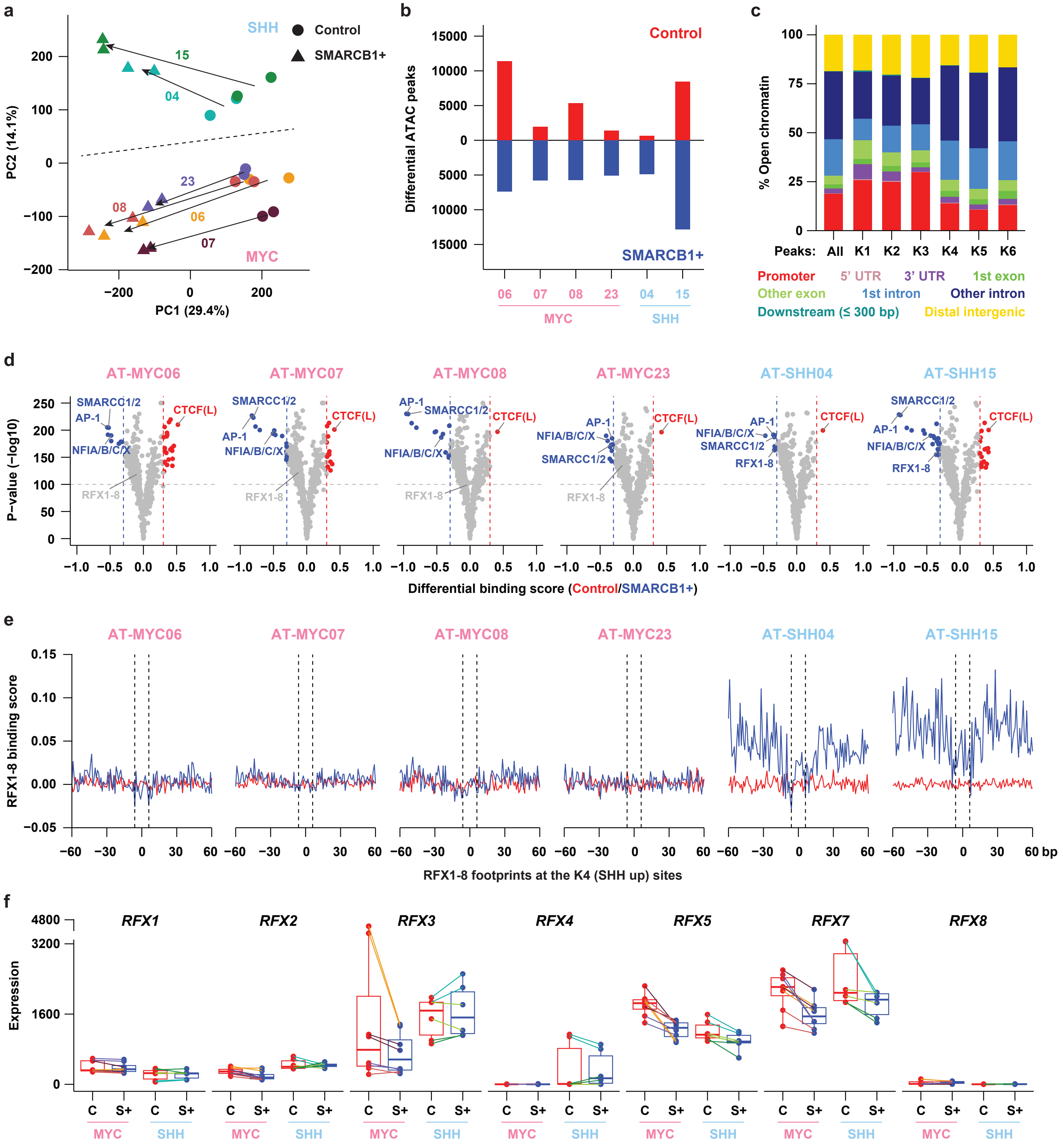

Extended Data Fig. 4

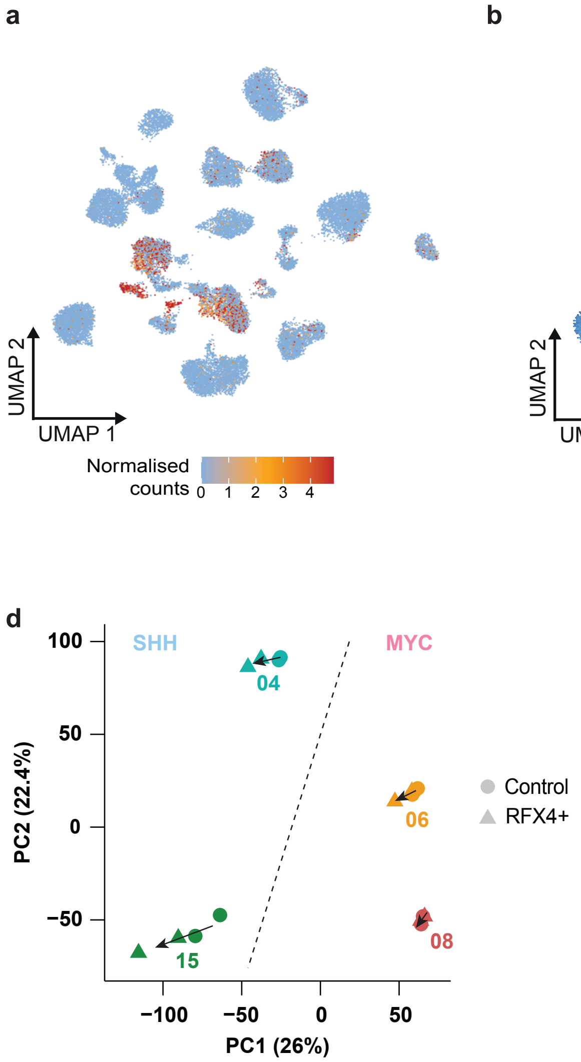

### Extended Data Fig. 5

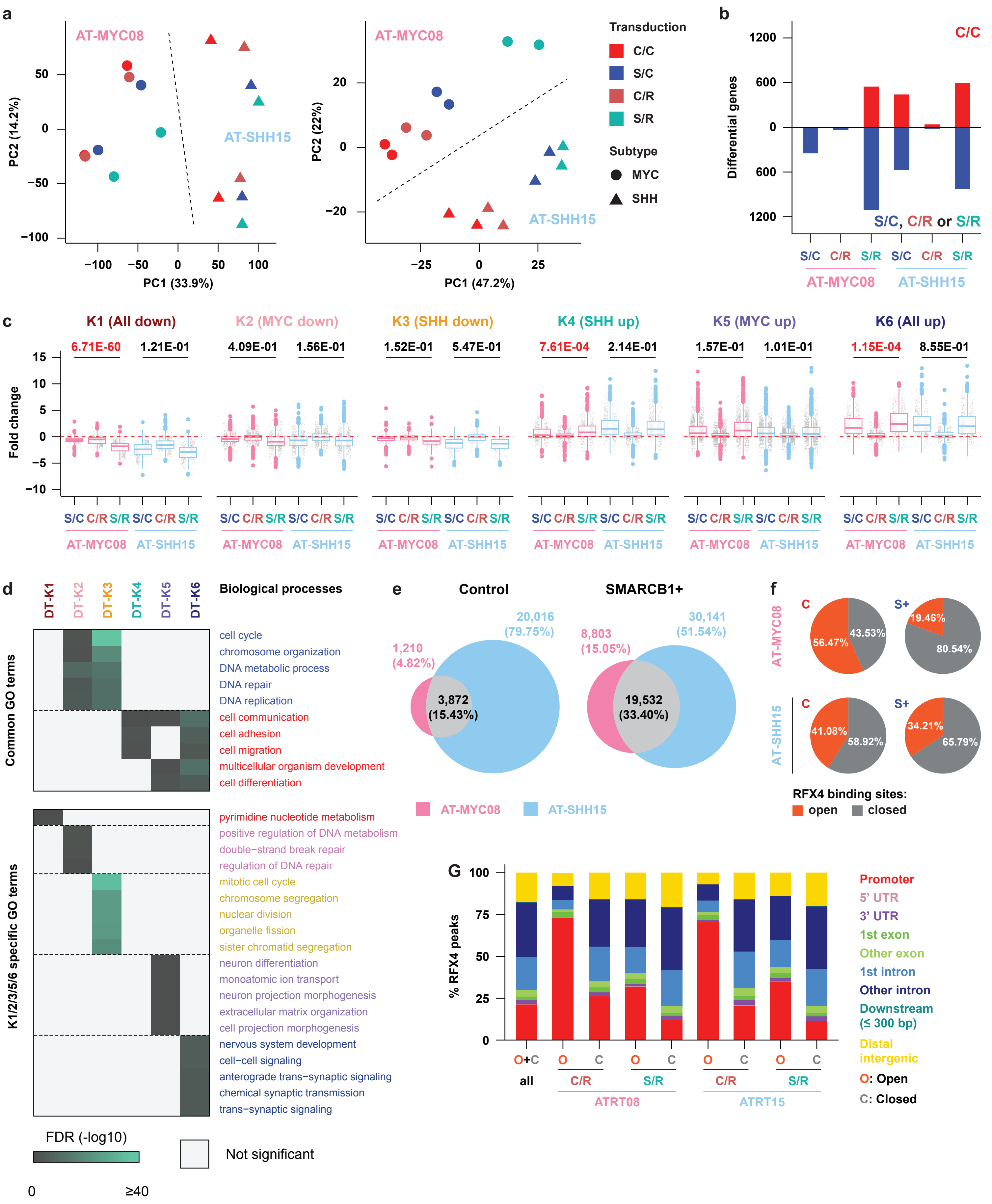

### Extended Data Fig. 6

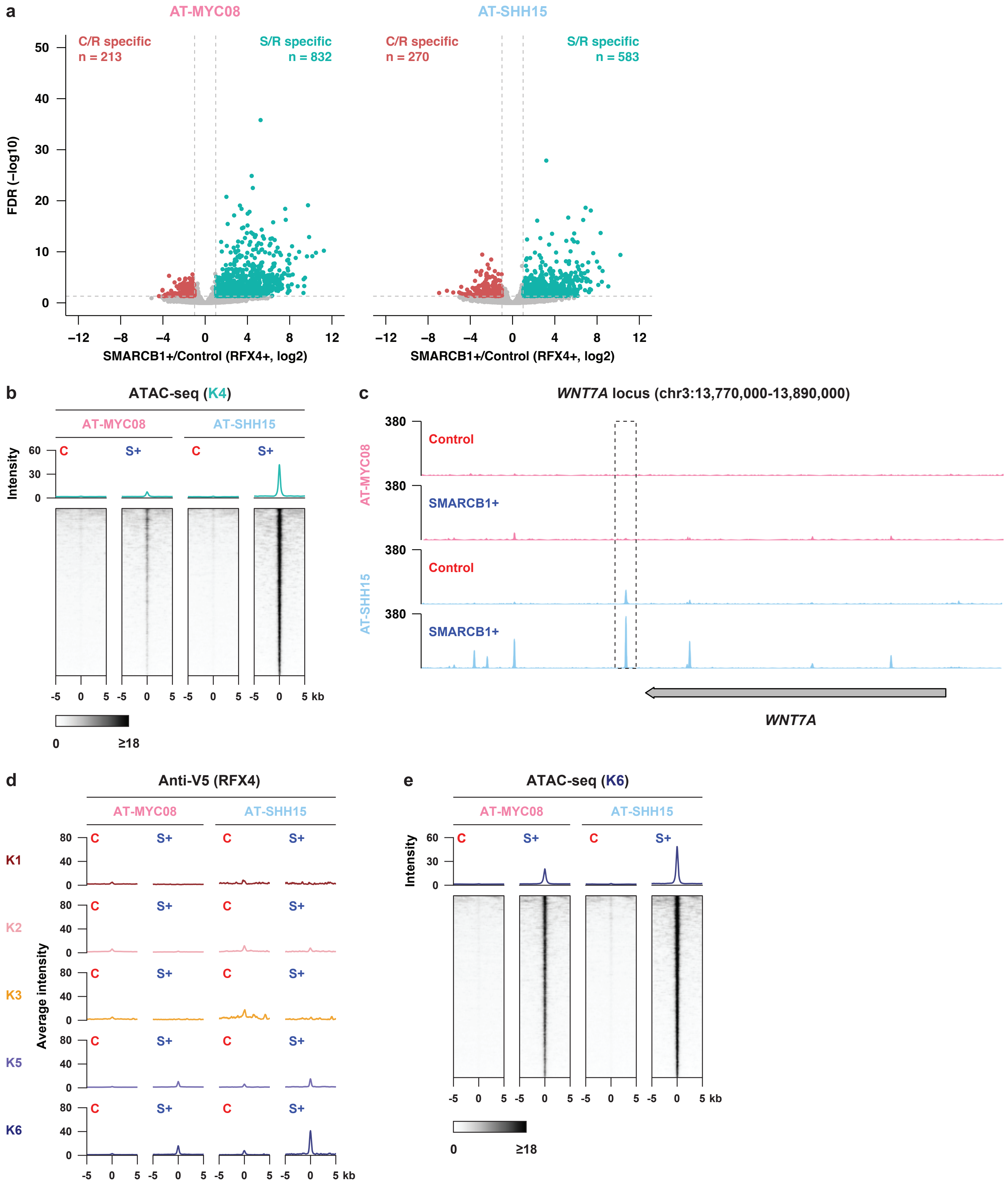
